# Patients with mesenchymal tumours and high *Fusobacteriales* prevalence have worse prognosis in colorectal cancer (CRC)

**DOI:** 10.1101/2021.05.17.444326

**Authors:** Manuela Salvucci, Nyree Crawford, Katie Stott, Susan Bullman, Daniel B. Longley, Jochen H.M. Prehn

## Abstract

**Objective:** Transcriptomic-based subtyping, Consensus Molecular Subtyping (CMS) and CRC Intrinsic Subtyping (CRIS), identify a patient subpopulation with mesenchymal traits (CMS4/CRIS-B) and poorer outcome. Here, we investigated the relationship between prevalence of *Fusobacterium nucleatum* (*Fn*) and *Fusobacteriales*, CMS/CRIS subtyping, cell type composition, immune infiltrates and host contexture to refine patients stratification and identify druggable context-specific vulnerabilities.

**Design:** We coupled cell culture experiments with characterization of *Fn*/*Fusobacteriales* prevalence and host biology/microenviroment in tumours from 2 independent CRC patient cohorts (Taxonomy: n=140; TCGA-COAD-READ: n=605).

**Results:** *In vitro, Fn* infection induced inflammation via NFκB/TNFα in HCT116 and HT29 cancer cell lines. In patients, high *Fn*/*Fusobacteriales* were found in CMS1, MSI tumours, with infiltration of macrophages M1, reduced macrophages M2, and high IL6/IL8/IL1β signaling. Analysis of the Taxonomy cohort suggested that *Fn* was prognostic for CMS4/CRIS-B patients, despite having lower *Fn* load than CMS1 patients. In the TCGA-COAD-READ cohort, we likewise identified a differential association between *Fusobacteriales* relative abundance and outcome when stratifying patients in mesenchymal (either CMS4 and/or CRIS-B) vs. non-mesenchymal (neither CMS4 nor CRIS-B). Patients with mesenchymal tumours and high *Fusobacteriales* had approximately 2-fold higher risk of worse outcome. These associations were null in non-mesenchymal patients. Modelling the 3-way association between *Fusobacteriales* prevalence, molecular subtyping, and host contexture with logistic models with an interaction term disentangled the pathogen/host-signaling relationship and identified aberrations (including EMT/WNT/NOTCH) as candidate targets.

**Conclusion:** This study identifies CMS4/CRIS-B patients with high *Fn*/*Fusobacteriales* prevalence as a high-risk subpopulation that may benefit from therapeutics targeting mesenchymal biology.

**Significance of this study:** *What is already known on this subject?:*

- *Fusobacterium nucleatum* (*Fn*), a commensal Gram-negative anaerobe from the *Fusobacteriales* order, is an onco-bacterium in CRC as a causal relationship between *Fn* prevalence and CRC pathogenesis, progression and treatment response has been reported *in vivo*.
- Broad spectrum antibiotics has proven moderately successful in reducing tumour growth in preclinical models. However, the use of antibiotics to treat bacterium-positive cases in the clinic is not a viable option as it may further alter the already dysbiotic gut microbiome of CRC patients and may also have limited efficacy against *Fn* which penetrates and embeds deeply within the tumour.
- The highly heterogenous CRC patient population can be classified into distinct molecular subtypes (CMS and CRIS) based on gene expression profiles mirroring the underlying transcriptional programs. Patients classified as CMS4 and CRIS-B exhibit a mesenchymal phenotype and have poorer outcome.

*What are the new findings?:*

- *Fn*/*Fusobacteriales* prevalence is associated with immune involvement (decrease in macrophages M1 and increase in macrophages M2) and activation of specific signalling programs (inflammation, DNA damage, WNT, metastasis, proliferation, cell cycle) in the host tumours.
- The prevalence of bacteria from the *Fusobacteriales* order, largely driven by *Fn* species, play an active or opportunistic role depending on the underlying host tumour biology and microenvironment.
- *Fn* and other species of the *Fusobacteriales* order are enriched in CMS1 (immuno, microsatellite unstable) patients compared to CMS2-4 cases.
- *Fn*/*Fusobacteriales* prevalence is associated with worse clinical outcome in patients with mesenchymal-rich CMS4/CRIS-B tumours, but not in patients with other molecular subtypes.

*How might it impact on clinical practice in the foreseeable future?:*

- *Fn*/*Fusobacteriales* screening and transcriptomic-based molecular subtyping should be considered to identify patients with mesenchymal-rich tumours and high bacterium prevalence and to inform disease management.
- *Fn*/*Fusobacteriales* prevalence may need to be addressed exclusively in patients with mesenchymal-rich high-stromal infiltrating tumours rather than a blanket-approach to treat all pathogen-positive patients.
- Clinical management of the disease for this subpopulation of high-risk patients with unfavourable clinical outcome could be attained by administering compounds currently in clinical trials that target aberrations in the host signaling pathways (NOTCH, WNT, EMT) and tumour microenviroment (inflammasome, activated T cells, complement system, and macrophage chemotactism and activation).

## Introduction

Colorectal cancer (CRC) has one of the highest morbidities and mortality rates among solid cancers and its incidence is steadily on the rise accounting for circa 10% of newly diagnosed cancer cases worldwide [1]. CRC patients with similar macroscopic clinico-pathological characteristics exhibit a high degree of heterogeneity at the molecular level, which translates into heterogeneous and often sub-optimal response to treatment. Thus, research has focussed on molecular subtyping strategies based on single or multi-omics data from the host to categorise patients into subgroups to aid in risk stratification and disease management. Subtyping strategies such as the Consensus Molecular Subtyping (CMS, [2]) and the Colorectal Cancer Intrinsic Subtyping (CRIS, [3]) classify patients into subgroups with more homogeneous signaling features based on key transcriptomic programs. Among the four subtypes identified by the CMS classifier, CMS4 patients have high stroma infiltration along with up-regulated angiogenesis and Transforming Growth Factor-β (TGFβ) signaling and show poorer recurrence-free and overall survival [2]. Similarly, CRIS-B patients feature mesenchymal traits and also exhibit poorer outcome compared to patients classified as CRIS-A, CRIS-C-E [3].

Recent research has identified the microbiome as a key player in health and disease, including cancer [4]. Several research groups, including ours, have shown that *Fusobacteriales*, largely from *Fusobacterium nucleatum* (*Fn*), are more abundant in tumour tissue compared to matched adjacent mucosa [5] suggesting a causative role in CRC progression [12]. More advanced, right-sided, MSI tumours are typically enriched with *Fn* [9]. Remarkably, anti-microbial treatment has been shown to reduce tumour burden in mouse xenograft models [10], corroborating the association between *Fn*-positive patients and poorer outcome observed in some studies [5]. However, the prognostic value of *Fn* prevalence was not observed in other cohort studies (reviewed in [16]). Thus, we hypothesized that the impact of *Fn*/*Fusobacteriales* may differ according to the underlying tumour biology.

In this study, we combined mechanistic *in vitro* experiments in colon cancer cells with an in-depth analysis in 2 independent CRC patient cohorts and a systematic multi-*omic* characterization of cell signalling and tumour microenvironment in n=745 patients to investigate the interaction between the dysregulation induced by *Fusobacteriales*, including *Fn*, prevalence on the human host and conversely, the characteristics of the host microenvironment that allow pathogens to thrive. Here, we provide evidence that the prognostic value of *Fn*/*Fusobacteriales* strongly relates to the molecular subtype of the host tumour and is confined to subtypes showing mesenchymal involvement.

## Results

### *Fusobacterium nucleatum* infection induces inflammation mediated by TNFα and NFκB in CRC cellular cultures

Due to the presence of *Fusobacterium nucleatum* (*Fn*) in CRC tumour tissue [5], a causative role for this bacterium to exacerbate tumourigenesis has been put forward. Infection of colon cells with *Fn* has previously been shown to induce inflammation, activate NFκB signaling and increase expression of the pro-inflammatory cytokine tumour necrosis factor alpha (TNFα) [18], (**Fig. 1A**). Hence, we infected HCT116 and HT29 colon cancer cell lines cultures for 6 hours to assess epithelial cell response to increasing amounts of *Fn* (multiplicity of infection, MOI, bacteria-to-cancer-cells 10, 100 and 1000). We found that NFκB signaling was indeed activated upon infection with *Fn* in CRC cell lines, as evidenced through the degradation of IκBα (alpha nuclear factor of kappa light polypeptide gene enhancer in B cells inhibitor) (**Fig. 1B**), an increase in NFκB transcriptional activity (**Fig. 1C**) and, a marked increase in mRNA expression of the NFκB target gene, TNFα (**Fig. 1D**). Taken together, these results confirm that *Fn* co-culture with human colon cancer epithelial cells promotes a pro-inflammatory response.

**Figure 1.**
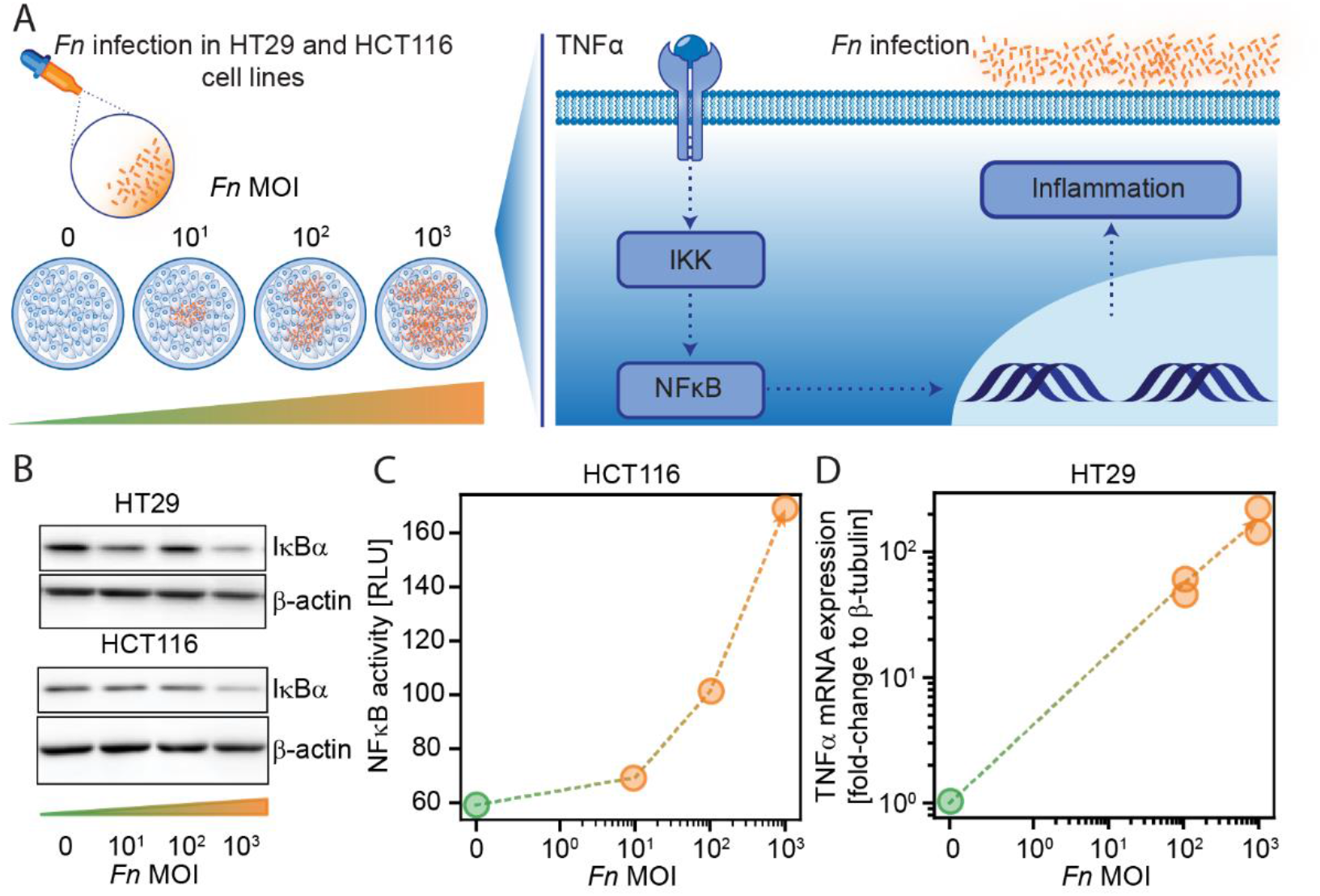
Fn infection induces inflammation mediated by TNFα and NFκB in HCT116 and HT29 CRC cell lines. **A**. Schematic representation of the experimental setup to investigate how *Fn* may trigger inflammation via TNFα and NFκB signalling pathways. **B**. Western blot analysis of IκBα and β-actin in HT29 and HCT116 cell cultures following infection with *Fn* (MOI bacteria-to-cancer-cells 10, 100 and 1000). **C**. NFκB transcriptional activity assay in HCT116 cells 6h following infection with *Fn* (MOI bacteria-to-cancer-cells 100 and 1000). **D**. TNFα mRNA expression relative to β-tubulin in HT29 cells 6h following infection with *Fn* (MOI bacteria-to-cancer-cells 100 and 1000). Panels **B-D** show representative results from duplicate experiments.

### Prevalence of *Fn* and *Fusobacteriales* in tumour resections

Next, we sought to investigate the relationship between inflammation in the human host and prevalence of *Fn* and *Fusobacteriales* in tumour resections of CRC patients. We selected an in-house multi-center stage II-III cohort (Taxonomy, n=140, [19], [20]) and the colon (COAD) and rectal (READ) cases of The Cancer Genome Atlas cohort (TCGA-COAD-READ, n=605 patients, **Fig. 2A**) to encompass the heterogeneity of the CRC clinico-pathological characteristics observed in the clinic. Demographic, clinico-pathological characteristics for the Taxonomy and TCGA-COAD-READ cohorts are summarised in **Suppl. Table 1**. We determined *Fn* abundance by a targeted quantitative real-time polymerase chain reaction (qPCR) in tumour resections of the Taxonomy cohort where we detected *Fn* in n=101 of 140 (72%) patients (**Fig. 2B**). The distribution of *Fn* positivity levels (relative to the human PGT gene) was heterogeneous and we categorized patients as *Fn*-high or *Fn*-low using the 75^th^ percentile as cut-off (**Fig. 2B**). We estimated *Fusobacteriales* relative abundance (RA) in the TCGA-COAD-READ cohort from RNA sequencing data by mapping non-human reads to microbial reference databases and retaining only high-quality matches (see **Methods**) with a PathSeq analysis [21], (**Fig. 2A**). For downstream analyses, we reported the relative abundance (RA) at the order, family, genus and species taxonomic rank and expressed it as percentage of the total bacterial abundance. We detected *Fusobacteriales* (defined as RA over zero, at the order level) in n=558 of 605 (92%) of the TCGA-COAD-READ patients, (**Fig. 2D**). *Fn* was the most abundant species and was detected in 82% of the TCGA-COAD-READ patients (compared to 72% in the Taxonomy cohort), accounting on average for approximately 45% of total *Fusobacteriales* RA and accounting for over 75% of total *Fusobacteriales* RA in 16% of cases (**Fig. 2C**). Analogously to the Taxonomy cohort, we categorized patients as *Fusobacteriales*-high or *Fusobacteriales*-low using the 75^th^ percentile as cut-off.

**Figure 2.**
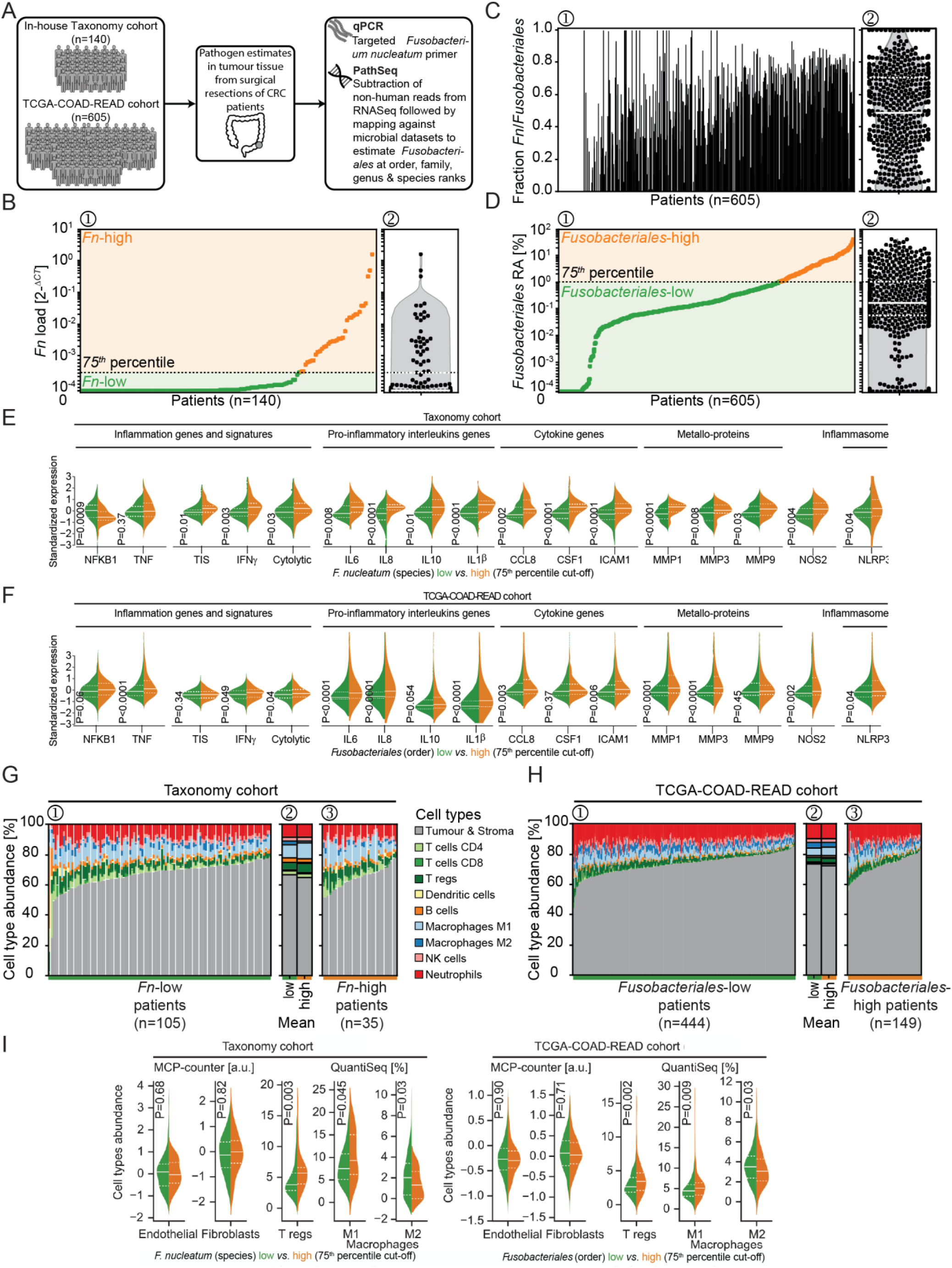
Fn/Fusobacteriales prevalence is associated with inflammation and immunosuppression in CRC patients of the Taxonomy and TCGA-COAD-READ CRC cohorts. **A**. Schematic representation of the cohorts included in the study and methods to estimate *Fn* load and *Fusobacteriales* (order) relative abundance in the Taxonomy and TCGA-COAD-READ cohorts, respectively. **B-D**. Per-patient (waterfall plot, **1**, left) and distribution (violin plot with overlaid data-points, **2**, right) of bacterium prevalence in tumour resections of the Taxonomy (n=140, **B**) and TCGA-COAD-READ (n=605, **D**). In **B-D 1**, patients are sorted in ascending order by prevalence of either *Fn* (Taxonomy cohort, **B**) or *Fusobacteriales* at the order taxonomic rank (TCGA-COAD-READ cohort, **D**). Cut-off of 75^th^ percentile used for patients’ stratification in downstream analysis is also indicated (black dotted line). Corresponding per-patient fraction of *Fn* species to total *Fusobacteriales* order relative abundance detected for the TCGA-COAD-READ cohort is shown in **C**. **E-F**. Violin plots grouped by prevalence of either *Fn* (Taxonomy cohort, **E**) or *Fusobacteriales* at the order taxonomic rank (TCGA-COAD-READ cohort, **F**) depicting the expression distribution of key genes or signatures involved in inflammation and immuno-suppression. Median and lower (25^th^) and upper (75^th^) percentiles are indicated by white solid or dashed lines, respectively. **G-H**. Stacked bar plots indicating cell type composition per-patient estimated from gene expression by quanTIseq in tumours with low vs. high prevalence of either *Fn* (Taxonomy cohort, **G**) or *Fusobacteriales* at the order taxonomic rank (TCGA-COAD-cohort, **H**). Cell type composition is shown sorted in ascending order of tumour and stromal content (**1** and **3**) and aggregated (by mean, **2** across the low- and high-subgroups). **I**. Distribution of specific tumour/stroma and immune cell types determined as indicated by either quanTIseq or MCPcounter grouped by either *Fn* (Taxonomy cohort) or *Fusobacteriales* at the order taxonomic rank (TCGA-COAD-READ cohort). Median and lower (25^th^) and upper (75^th^) percentiles are indicated by white solid or dashed lines, respectively.

### Higher *Fn* and *Fusobacteriales* prevalence correlates with inflammation and immune involvement

We examined the association between host gene expression profiles of key markers shown to orchestrate inflammation and either *Fn* load or *Fusobacteriales* RA in the Taxonomy and TCGA-COAD-READ cohorts, respectively. In line with the *in vitro* experiments (**Fig. 1**), we detected an increase in NFKB1 and a trend in TNFα gene expression, recapitulated by transcriptomic-based signatures for an overall inflammation status (TIS) mediated by the cytolytic and interferon (IFNγ) pathways in the Taxonomy cohort (**Fig 2E**). When investigating further key inflammation players, we observed a marked increase in pro-inflammatory interleukins (IL6, IL8, IL10, IL1β, IL13), cytokines/chemokines (CCL8, CSF1, ICAM1), metallo-proteins (MMP1, MMP3, MMP9), NOS2, the inflammasome complex (NLRP3) and COX2 in *Fn*-high vs. -low Taxonomy patients (**Fig. 2E** and **Suppl. Fig. 1**).

Next, we sought to validate and build upon our findings from the in-house Taxonomy cohort by analyzing the TCGA-COAD-READ cohort (**Fig. 2F**). At the transcription level, we confirmed an exacerbated inflammatory state when comparing *Fusobacteriales*-high and -low patients mediated by the NFκB-TNFα axis, IFNγ with cytolytic involvement. *Fusobacteriales*-high patients overexpressed pro-inflammatory interleukins (IL6, IL8, IL10, IL1β), cytokines/chemokines (CCL8, ICAM1), metallo-proteinases (MMP1, MMP3), NOS2 and inflammasome markers (NLRP3), (**Fig. 2F**).

As inflammation is strongly tied to immune cell migration and activity, we next investigated whether there was a link between immune cell composition and either *Fn* load (Taxonomy) or *Fusobacteriales* RA (TCGA-COAD-READ). Cell composition was computationally deconvoluted from gene expression profiles with quanTiseq [23] and MCP-Counter [24], (**Fig. 2G-H**). Despite observing high inter-patient heterogeneity in cell composition within the Taxonomy and TCGA-COAD-READ cohorts, we robustly detected higher immune cell activation and polarization when comparing patients with high vs. low *Fn* load (Taxonomy) or *Fusobacteriales* RA (TCGA-COAD-READ). Patients with high *Fn* load (Taxonomy) or *Fusobacteriales* (TCGA-COAD-READ) showed higher predicted abundance of regulatory T cells (T regs) coupled with an increase in M1 macrophages and decrease in M2 macrophages (**Fig. 2I**). MCP-counter identified a strong positive association between neutrophil infiltration and either *Fn* load (Taxonomy) or *Fusobacteriales* RA (TCGA-COAD-READ), (**Fig. 2I**). However, no difference in predicted neutrophils abundance was detected by quanTIseq. Importantly, no difference in fibroblasts and endothelial cells was observed by *Fn*/*Fusobacteriales* in either cohort by either method (**Fig. 2I**).

### Multi-*omic* characterization of the association between *Fusobacteriales* relative abundance and human host tumour microenvironment in the TCGA-COAD-READ cohort

We next leverage the rich molecular characterization of the TCGA-COAD-READ cohort to perform a systematic and unbiased characterization of the association between *Fusobacteriales* RA and patient clinical and molecular features to identify human host vulnerabilities that may be conducive for tumour development (**Fig. 3**).

**Figure 3.**
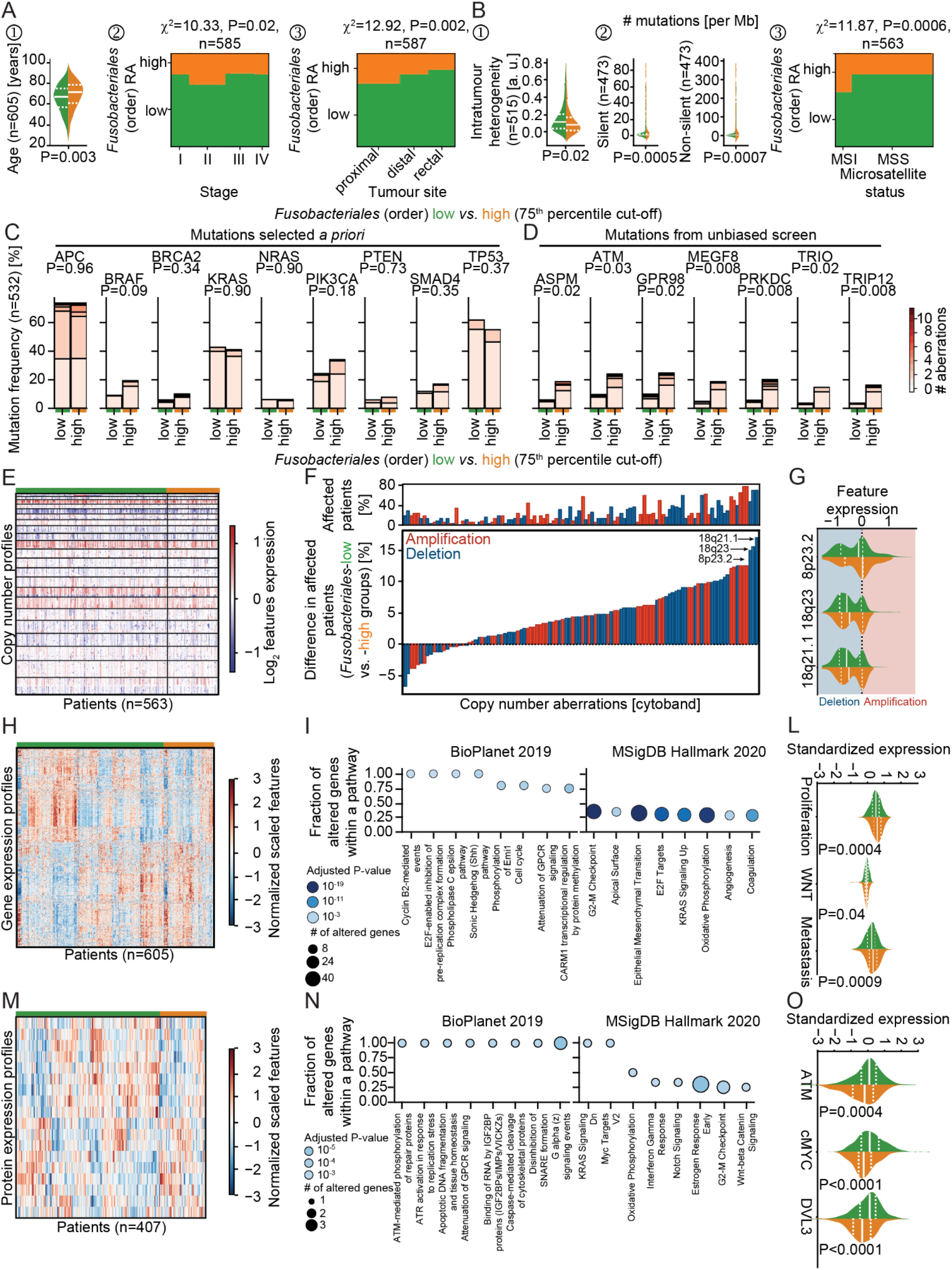
Multi-omic characterization of the association between Fusobacteriales relative abundance and human host tumour microenvironment in the TCGA-COAD-READ cohort. **A-B**. Association between *Fusobacteriales* at the order taxonomic rank binned into high vs. low (cut-off 75^th^ percentile) and clinico-pathological (**A**) and mutational (**B**) characteristics of the human host. **C-D**. Comparison of frequency of occurrence of mutations selected *a priori* (**C**) or identified by an unbiased scan (**D**) in *Fusobacteriales*-high vs -low patients. Colorbar indicates number of detected aberrations among frame shift deletions and insertions, in frame deletions and insertions, missense and nonsense mutations and splice sites. P-values were computed with χ^2^ independence tests and adjusted for multiple comparisons (Benjamini-Hochberg false discovery rate). **E-G**. Heatmap (**E**) displaying copy number alterations grouped by *Fusobacteriales*-high (in orange) and -low (in green) relative abundance. Waterfall plot (**F**) displaying differences in recurrent copy number aberrations detected in patients with low-vs. high *Fusobacteriales*. Top panel in **F** reports percentage of patients affected by recurrent copy number aberrations. Distribution of top 3 deletions whose frequency of occurrence differs between *Fusobacteriales*-high and -low patients (**G**). Red and blue shading indicates amplification and deletions, respectively. **H-L**. Heatmap (**H**) displaying expression of genes differentially expressed when comparing *Fusobacteriales*-high vs. low patients and corresponding pathway enrichment analysis (**I**). Expression distribution grouped by *Fusobacteriales* RA for selected gene expression signatures is shown in **L**. **M-O**. Heatmap (**M**) displaying expression of proteins differentially expressed when comparing *Fusobacteriales*-high vs. low patients and corresponding pathway enrichment analysis (**N**). Expression distribution grouped by *Fusobacteriales* RA for key proteins is shown in **O**. In violin plots, the median and lower (25^th^) and upper (75^th^) percentiles are indicated by white solid or dashed lines, respectively. Orange and green annotation bars denote patients with high vs. low *Fusobacteriales* relative abundance (75^th^ percentile cut-off). (Unadjusted) P-values in **L** and **O** were determined by Kruskal-Wallis H-test for independent samples.

We observed higher *Fusobacteriales* in patients of older age, diagnosed with more advanced disease stage and tumours located in the colon, particularly in proximal sites, (**Fig 3A**), corroborating studies assessing *Fn* [13]. In contrast, we found no statistically significant differences in *Fusobacteriales* RA by sex, body mass index and neither lymphovascular nor perineural invasion (**Sup. Fig. 2**).

Patients harbouring higher *Fusobacteriales* showed lower genomic intra-tumour heterogeneity, had higher silent and non-silent mutational burden and were enriched in microsatellite unstable cases, (**Fig 3B**). *Fusobacteriales*-high patients showed an increase in transitions, defined as the exchange of two-ring purines (A↔G) or of a one-ring pyrimidines (C↔T), coupled with a decrease in transversions, a substitution of purine for pyrimidine bases (**Suppl. Fig. 3A**) as evidenced by a decrease in conversion changes of C>G and T>A, (**Suppl. Fig. 3B**). We found no difference in prevalence of common mutations in CRC by *Fusobacteriales* (high vs. low) except for BRAF (**Fig. 3C**). BRAF mutations trended to be more common among *Fusobacteriales*-high patients, as observed when assessing *Fn* [13]. A comprehensive screen revealed that mutations in cell cycle (ATM), Hedgehog signaling (MEGF8), DNA damage/repair (TRIP12, PRKDC), mitotic spindle (ASPM), migration/adhesion (TRIO, GPR98) were more prevalent in *Fusobacteriales*-high patients, (**Fig. 3D, Suppl. Table 2**).

Next, we set out to investigate the relationship between copy number alterations (CNAs) and *Fusobacteriales* presence in the TCGA-COAD-READ cohort (**Fig. 3E-G**). We determined recurrent CNAs amplifications and deletions across the whole cohort by applying the GISTIC algorithm [26] (**Sup. Figs. 4-5** and **Suppl. Table 3**). *Fusobacteriales*-high cases showed lower chromosomal instability with a lower fraction of the genome affected by recurrent CNAs, in line with their MSI unstable status. Next, we identified CNA amplifications (in red) or deletions (in blue) whose frequency of occurrence differed when comparing *Fusobacteriales*-high vs. -low patients and thus may be specifically associated with the bacterium presence (**Fig. 3F**). CNAs more frequently (>15%) observed in *Fusobacteriales*-high vs. low cases included deletions in 8p23.2 (tumour suppressor CSMD1 and LOC100287015); 18q21.1 (MIR4743 and RNA binding by CTIF) and 18q23 which impacts the regulation of interleukin-6 and chemokine secretion, cell-cell adhesion and host of viral transcription, as determined by enrichment analyses carried out with EnrichR, (**Fig. 3G**).

**Figure 4.**
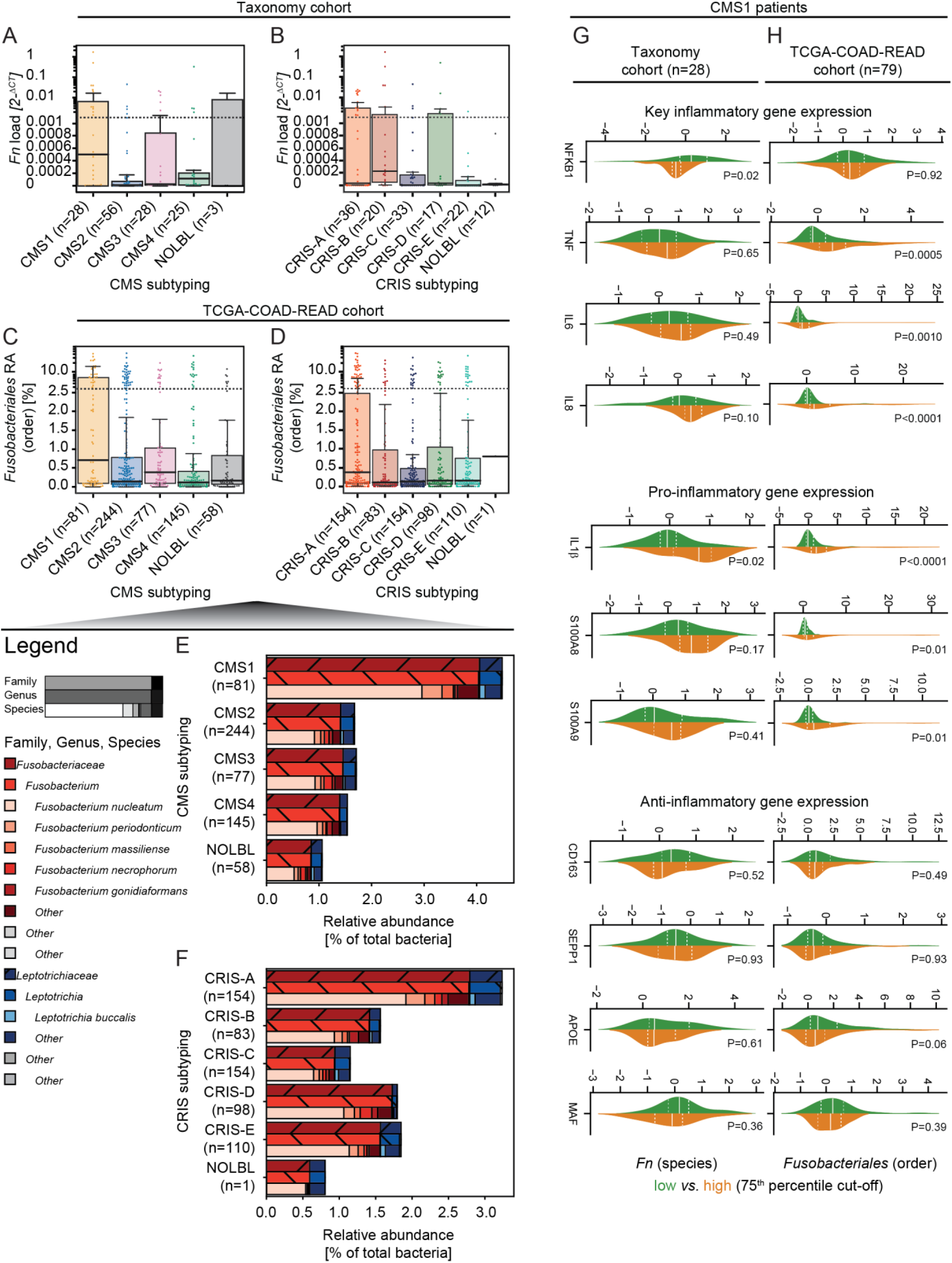
Prevalence of Fn/Fusobacteriales by transcriptomic-based molecular subtypes of the host. **A-D**. Boxplot with overlaid dot plots displaying the dependency by CMS (**A, C**) and CRIS (**B, D**) molecular subtyping by prevalence of either *Fn* (Taxonomy cohort, **A-B**) or *Fusobacteriales* at the order taxonomic rank (TCGA-COAD-READ cohort, **C-D**). **E-F**. Relative abundance (to total bacterial kingdom) of *Fusobacteriales* reported at increasing resolution of taxonomic rank (family, genus and species) by CMS (**E**) and CRIS (**F**) subtypes (aggregated by mean). **G-H** Distribution of key (pro-)/(anti-)inflammatory genes grouped by either *Fn* (Taxonomy cohort, **G**) or *Fusobacteriales* at the order taxonomic rank (TCGA-COAD-READ cohort, **H**) restricted to CMS1 patients. Median and lower (25^th^) and upper (75^th^) percentiles are indicated by white solid or dashed lines, respectively. (Unadjusted) P-values were determined by Kruskal-Wallis H-test for independent samples.

We then focused on the transcriptional level and we combined enrichment analyses with pathway-activity signatures to compare cellular processes by *Fusobacteriales* RA (**Fig. 3H-L**). Transcriptional profiles differed by mTORC1 and Myc signalling, cell cycle (G2-M checkpoint), mitotic spindle, epithelial-to-mesenchymal transition, TGFβ and interleukin-1 regulation of extracellular matrix, matrix remodelling including focal adhesion, cytoskeleton and contractile actin filament bundle, mitochondrial translational elongation/termination and protein complex assembly and stromal estimates (**Fig. 3H-I, Sup. Fig. 6** and **Sup. Table 4**). We corroborated these findings by comparing the activation of signalling pathways estimated by gene set signatures identified in the literature (see **Methods**) in *Fusobacteriales*-high vs. low patients. Indeed, *Fusobacteriales* presence was positively associated with proliferation, WNT, metastasis (**Fig. 3L**) and DNA damage.

Next, we sought to investigate whether the findings at the genomic and transcriptional level were also observed in protein profiles determined by Reverse Phase Protein Array (RPPA). We found a differential expression by *Fusobacteriales* RA for proteins involved in microenvironment composition (Claudin7), cell cycle (Cycline1), apoptosis (cleaved Caspase7), proliferation (DLV3), Hippo pathway (Yap), DNA damage (Chk1, ATM), receptor and MAP kinases and PI3K signalling, (**Fig. 3M-O, Sup. Fig. 7** and **Sup. Table 5**).

### *Fn* and *Fusobacteriales* prevalence differs by transcriptomic-based molecular subtype

The systematic screen above pinpointed host aberrations by *Fusobacteriales* prevalence that are hallmarked by transcripomic-based molecular subtypes. Hence, we classified patients in the study by CMS [2] and CRIS [3] subtyping. We observed higher *Fn* load (Taxonomy, **Fig. 4A**) and *Fusobacteriales* RA (TCGA-COAD-READ, **Fig. 4C**) in CMS1 tumours, corroborating the interplay between pathogen prevalence and host immunity. Moreover, we observed higher *Fn* load in CRIS-B tumours (**Fig. 4B**) and *Fusobacteriales* RA in CRIS-A cases (**Fig. 4D**) of the Taxonomy and TCGA-COAD-READ cohorts, respectively. At the family rank, *Fusobacteriaceae* were more abundant than *Leptotrichiaceae* accounting for 77% and 23% of total *Fusobacteriales* RA and ∼2% and ∼<1% of the total bacteria RA, respectively. In line with the findings at the order level, we observed an increase in *Fn*, the most abundant *Fusobacterium* species, in CMS1 and CRIS-A cases (**Fig. 4E-F**). In line with the findings at the order level, we observed an approximately 3-fold increase when comparing patients classified as CMS1 vs. the rest (**Fig 4E**). *Fn*, the most abundant *Fusobacterium* species, was enriched in CMS1 and CRIS-A cases (**Fig. 4E-F**). Next, we examined whether the positive association between inflammation and immune involvement by *Fn*/*Fusobacteriales* presence could be ascribed to the host CMS1 milieu or whether there was an additional pathogen-induced component. When restricting the analysis to CMS1 cases, we observed higher expression of pro-inflammatory markers in *Fusobacteriales*-high patients of the TCGA-COAD-READ cohort. We detected no association between pathogen prevalence and expression of anti-inflammatory markers or inflammation signatures in neither CRC cohorts (**Fig. 4G-H**). Taken together these results suggest that *Fn*/*Fusobacteriales* may play an active role in mediating inflammation in the host.

### Patients with high *Fn*/*Fusobacteriales* have worse outcome in CMS4/CRIS-B

Next, we sought to investigate whether bacterium presence correlated with patient clinical outcome assessed by overall-(OS), disease-specific-(DSS) and disease-free-(DFS) survival endpoints (**Fig. 5** and **Suppl. Figs. 8-10**).

**Figure 5.**
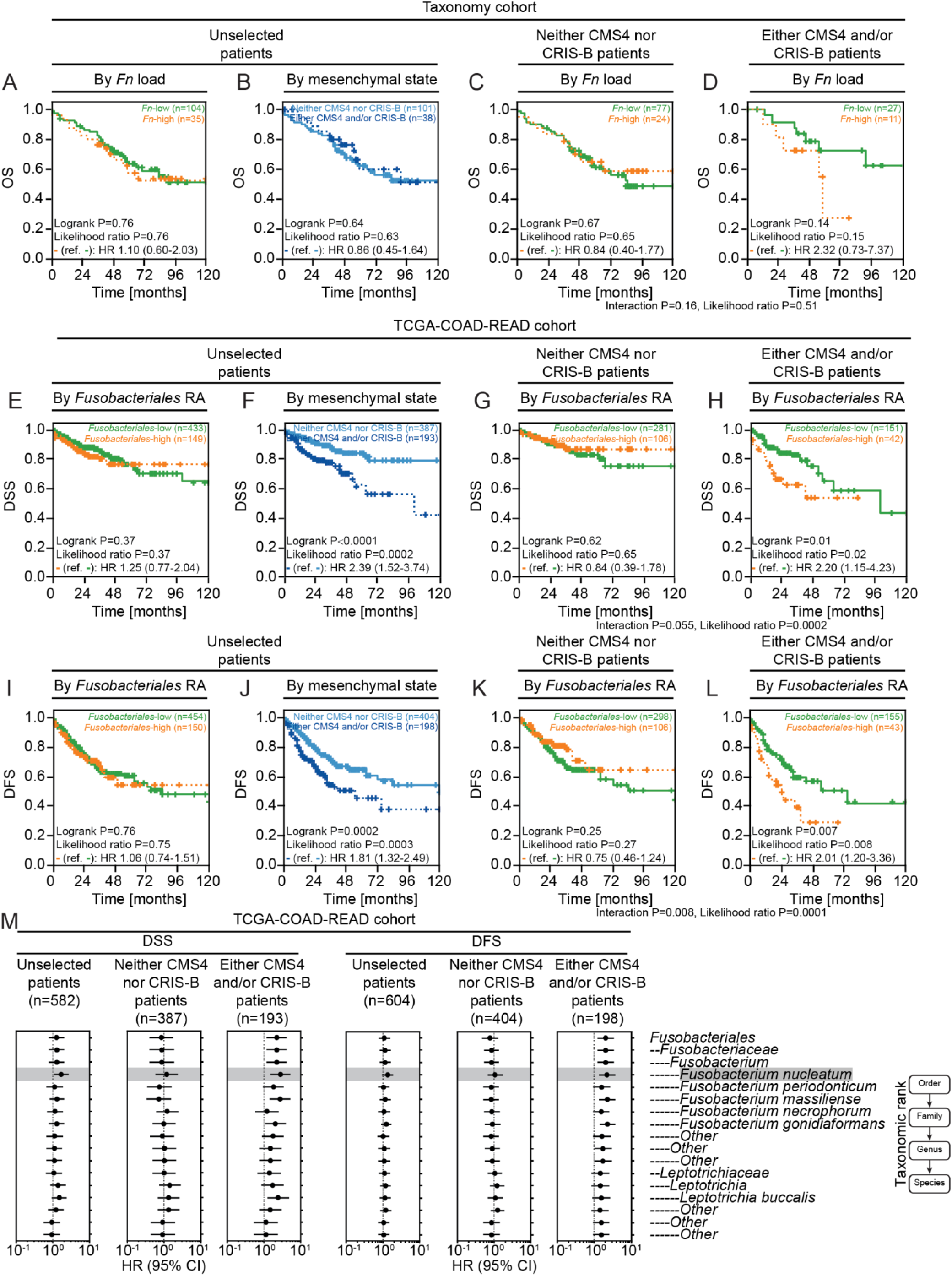
High Fn/Fusobacteriales prevalence is associated with negative clinical outcome in patients with mesenchymal-like tumours. **A-L**. Kaplan-Meier estimates comparing survival curves in patients of the Taxonomy (OS, **A-D**) and TCGA-COAD-READ (DSS and DFS, **E-L**) cohorts. Patients across the whole cohort were grouped by prevalence (high vs. low based on 75^th^ percentile cut-off) in **A, E, I** or mesenchymal status (CMS4 and/or CRIS-B vs. remaining cases) in **B, F, J**. Patients were grouped by prevalence and further stratified by mesenchymal status in **C-D, G-H, K-L**. Prevalence refers to either *Fn* load or *Fusobacteriales* RA at the order level for the Taxonomy and TCGA-COAD-READ cohorts, respectively. **M**. Cox regression models fitted on bacterium RA reported at the order, family, genus and species taxonomic ranks. For each taxonomic rank, patients were classified as low or high prevalence using the corresponding 75^th^ percentile RA abundance as cut-off. Univariate Cox regression models were fitted when evaluating association between pathogen prevalence (high vs. low; reference low) at each taxonomic rank and either DSS or DFS in the whole unselected patient population (left panel). Cox regression models with an interaction term between pathogen prevalence (high vs. low; reference low) and mesenchymal status (mesenchymal, i.e. either CMS4 and/or CRIS-B, vs. non-mesenchymal, i. e. neither CMS4 nor CRIS-B) at each taxonomic rank and either DSS or DFS were fitted to evaluate differential impact of bacterium on clinical outcome by tumour biology (right panels).

**Figure 6.**
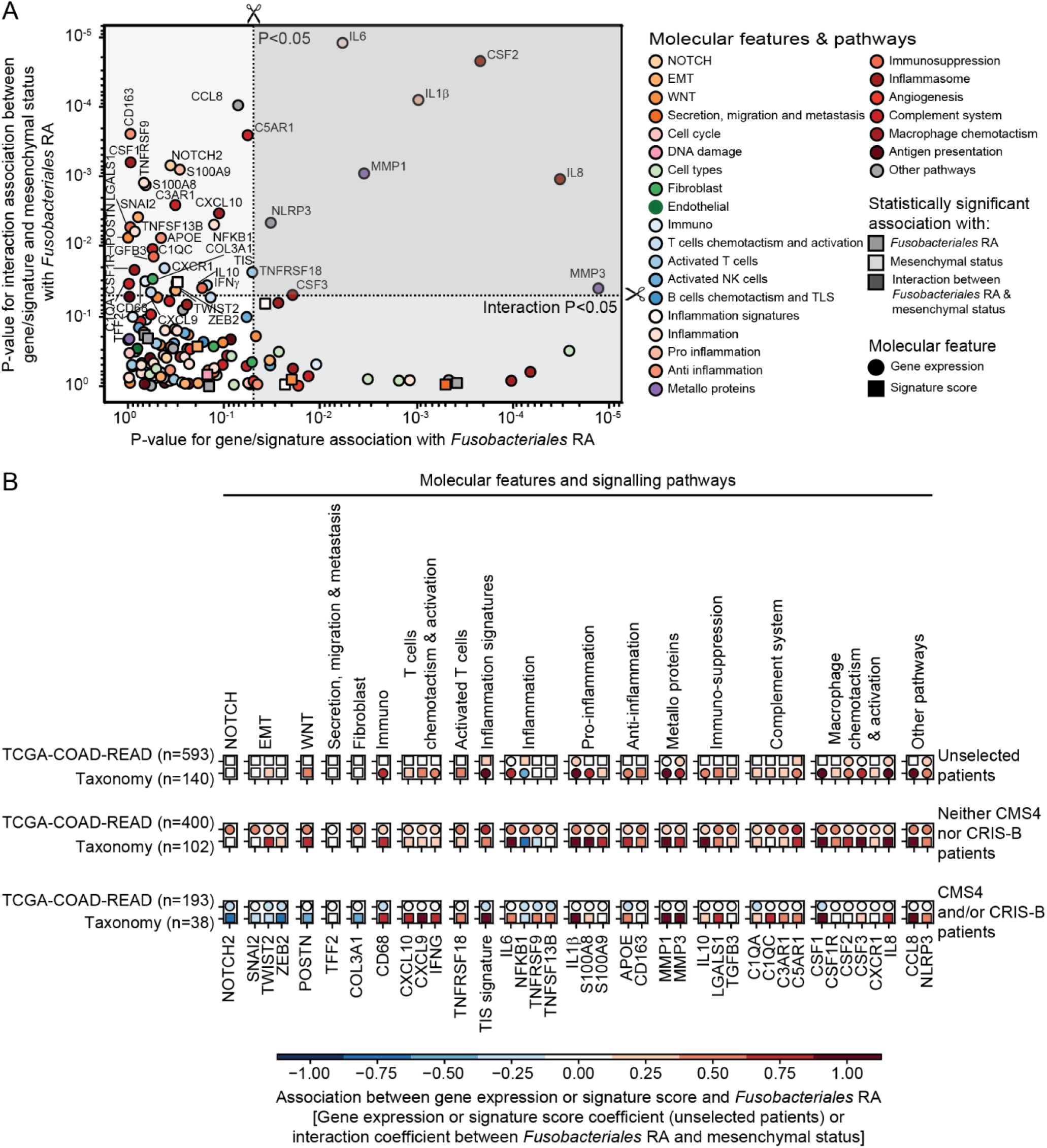
Exploration of mechanism underlying differential impact of Fusobacteriales in mesenchymal vs. non-mesenchymal tumours. **A**. Scatterplot depicting P-values derived by assessing with logistic regression models the relationship between genes/signatures associated with *Fusobacteriales* RA in univariate analysis (model 1, x axis) or the interaction with mesenchymal status (model 2, y axis). **B**. Breakdown of association including direction, effect size, in the unselected patients’ population and within mesenchymal vs. non-mesenchymal cases. Only gene/signatures with significant interaction between *Fusobacteriales* RA and the gene/signature interaction with the molecular subtype (model 2) in the TCGA-COAD-READ cohort are included. Associations for both the TCGA-COAD-READ (*Fusobacteriales* RA) and Taxonomy (*Fn* load) cohorts are shown. Statistically significant associations are represented with circle markers whereas non-significant associations are indicated by squared markers.

We found no statistically significant differences in neither cohort when comparing survival curves from patients grouped by either *Fn* load or *Fusobacteriales* RA (**Fig. 5A, E, I** and **Suppl. Figs. 8-10**). We hypothesized that *Fn*/*Fusobacteriales* may result in poorer outcome in a subtype-dependent context. Indeed, we identified a differential association between *Fusobacteriales* RA and clinical outcome of the TCGA-COAD-READ cohort in mesenchymal (either CMS4 and/or CRIS-B) vs. non-mesenchymal (neither CMS4 nor CRIS-B) tumours, (**Fig. 5G, H, K, L** and **Sup. Figs. 8-10**). *Fusobacteriales*-high mesenchymal patients had approximately 2-fold higher risk of worse outcome while these associations were null in non-mesenchymal patients (**Fig. 5G, H, K, L** and **Sup. Figs. 8-10**).

Although numbers in the Taxonomy cohort are limited, when restricting the analysis to CMS4 and/or CRIS-B cases, we observed a trend whereby *Fn*-high patients had shorter OS than those with low *Fn* load. Again, no difference in survival curves by *Fn* load was observed in non-mesenchymal Taxonomy patients (**Fig. 5C-D**).

Exploratory analyses examining the association between clinical outcome and pathogen prevalence at taxonomic ranks of increasing resolution (order, family, genus and species) in the TCGA-COAD-READ cohort by fitting Cox regression models on the whole unselected population and in mesenchymal vs. non-mesenchymal settings revealed that the prognostic impact stems primarily from, but is not limited to, species, including *Fn*, from the *Fusobacterium* genus from the *Fusobacteriaceae* family (**Fig. 5M** and **Sup. Fig. 10**).

### Putative mechanisms underlying selective *Fusobacteriales* virulence in mesenchymal tumours

We examined the host signaling pathways and microenvironment to identify alterations that may be mediated by and/or exacerbated by *Fusobacteriales* (i.e. interact) and, thus, may promote virulence and, ultimately, result in an unfavorable clinical outcome. To disentangle the 3-way association between *Fusobacteriales* RA, gene/signature, and molecular subtyping, we fitted 2 distinct logistic regression models for each feature of interest in the TCGA-COAD-READ cohort. The selection of features was hypothesis-driven and included key host signaling pathways and immuno-modulators (**Fig. 6A**).

**Fig. 6A** reports adjusted P-values from the 2 models capturing the association between *Fusobacteriales* RA (high vs. low) and either each gene/signature (**model 1**: *Fusobacteriales ∼ gene/signature*, x-axis) or the interaction between each gene/signature with the molecular subtype (**model 2**: *Fusobacteriales ∼ gene/signature:molecular subtype*, y-axis). The top right half quadrant (darker gray shaded area) identifies a set of genes/signatures whose expression patterns differ by molecular subtype (statistically significant interaction p-value in model 2) and thus may be mediating the pathogenic impact of *Fusobacteriales* and were prioritized for downstream analyses (**Fig. 6B**).

NOTCH, EMT, TIS score, IL6, CSF1 are among the genes/signatures identified by model 2 in **Fig. 6A** whose expression profiles track with molecular subtyping and may represent druggable vulnerabilities in patients with mesenchymal tumours and high *Fn*/*Fusobacteriales* prevalence and ameliorate clinical outcome (**Fig. 6B**).

## Discussion

*Fusobacteriales*, predominantly *Fn*, have been associated [5] with pathogenesis, progression and treatment response in CRC. We coupled mechanistic studies in cell cultures with hypothesis-driven and unbiased screening in clinically-relevant and ‘omics-rich CRC cohorts to examine the cross-talk between pathogen-host and pathogen-tumour microenvironment. We demonstrated relationships between *Fn*/*Fusobacteriales* prevalence with host immune, signaling activation and transcriptomic-based molecular subtypes. Our findings suggest that host-pathogen interactions can define patient sub-populations where *Fn*/*Fusobacteriales* play an active or opportunistic role depending on the underlying host tumour biology and microenvironment and identify putative druggable and clinically-actionable vulnerabilities.

We observed higher *Fn*/*Fusobacteriales* prevalence in CMS1 patients, corroborating findings by Purcell [28]. Interestingly, we found that higher pathogen prevalence did not correlate with poorer disease outcome. In contrast, *Fn*/*Fusobacteriales* virulence was exacerbated in CMS4/CRIS-B patients, suggesting that pathogen persistence may need addressing exclusively in mesenchymal-rich high-stromal infiltrating tumours and arguing against a blanket-approach to treat all pathogen positive patients. Treatment with wide spectrum antibiotics reduces the growth of *Fn*-positive tumours *in vivo* [10]. However, the use of antibiotics to treat *Fn*-positive CRC tumours may be limited as *Fn* penetrate deeply within tumour, immune and endothelial cells where they internalize with endosomes and lysosomes [29], adapt [30] and persist [10]. In addition, long-term use of antibiotics can cause dysbiosis.

Given that “it takes two to tango”, namely a high pathogen prevalence and a conducive host milieu, we further examined this interdependence to identify druggable aberrations in the host signaling pathways and microenvironment. We identified putative targets related to (pro-)inflammation, inflammasome, activated T cells, complement system, metallo-proteins and macrophage chemotaxis and activation. *Fusobacteriales* induce a constitutively activated NF-kB-TNFα-IL6 state which results in activation of metallo proteins and inflammatory cytokines (CSF1-3) which mediate macrophage differentiation, inhibit cytotoxic immune cells and promote proliferation of myeloid-derived-suppressor (MDSC) cells. Indeed, we observed an increase in inflammation and macrophages M1 and decrease in macrophages M2 in patients with higher *Fn*/*Fusobacteriales* prevalence. We envisage that therapeutic options, such as NLRP3/AIM2 inflammasome suppression [31], IL1β blockade [32], TNFα [33] or IL6 inhibition [34], that have been approved for treatment of chronic inflammation and cytokines storm syndrome in multiple cancers, rheumatoid arthritis and COVID-19 may ameliorate the immunosuppressive microenvironment induced by *Fn*/*Fusobacteriales*.

Importantly, these targets are involved in not only promoting an immunosuppressive microenvironment by recruiting tissue-associated macrophages (TAMs) and MDSCs, but also in orchestrating invasion, angiogenesis, epithelial-to-mesenchymal transition and, ultimately, metastasis. The pro-metastatic role of *Fn*/*Fusobacteriales* is further corroborated by findings in the literature linking higher pathogen prevalence in more-advanced disease stage and metastasis in clinical specimens [5] and higher metastatic burden in mice inoculated with *Fn* [35].

Cancer cells with an EMT phenotype secrete cytokines such as IL10 and TGFβ that can further promote an immunosuppressive microenvironment. Additionally, secretion of IL6 and IL8 from stroma cells can further foster an EMT phenotype, activate primary fibroblasts (carcinoma-associated fibroblast, CAFs) which, in turn, may promote angiogenesis and invasion [36]. Taken together, these aberrations may result in a self-reinforcing mechanism that confers on cancer cells the ability to migrate, invade the extracellular matrix, extravasate and seed metastasis. Indeed, when comparing the transcriptomic profiles by *Fusobacteriales* RA in the TCGA-COAD-READ cohort, we identified dysregulation affecting cell architecture involving apical surface dynamics and Aurora A kinase signaling, which regulate cMyc, DNA repair, cell motility/migration and induce EMT transition via β-catenin and TGFβ leading to metastasis and resistance to treatment in multiple cancer types [37]. Small molecule inhibitors against aurora A have shown encouraging results in preclinical studies and clinical trials in CRC [38] and other cancers [39]. Cytoskeleton shape, filopodium protrusions and alterations in cell adhesion and structure are hallmark of extracellular matrix invasion. EMT key effectors, SNAIL and ZEB1, alter apical surface dynamics by inhibiting scaffolding proteins and by inducing expression of matrix metalloproteins (MMP3, MMP9), resulting in loosened tight-junctions, altered cell polarity and increased plasticity which, in turn, enable cell invasion [40]. Dysregulations in MMPs expression may aid cancer cells that have reached the bloodstream to extravasate to distant tissues [41] by priming the vascular endothelium via upregulation of VEGF-A [42] and by increasing permeability via COX2 upregulation [43]. Our analyses in the TCGA-COAD-READ cohort identified higher expression of VEGF as well as an angiogenesis signature and COX2 in patients with higher *Fusobacteriales* RA. MMPs treatment with a new generation of selective and highly penetrative inhibitors [44] is being trialed in gastrointestinal cancers [45] and Mehta reported lower *Fusobacteriales* RA in subjects treated with Aspirin, a COX2 inhibitor [46].

Green [47] demonstrated that MAPK7 is a master regulator of MMP9 and promotes the formation of metastasis. Indeed, we observed a dysregulation in MAPK signaling at the protein level when comparing *Fusobacteriales*-high vs. -low patients of the TCGA-COAD-READ cohort. MAPK7 induces EMT transition, cell migration and regulates TAMs polarization in a metallo proteins-dependent manner [47], rendering it an appealing upstream therapeutic target. IL6 orchestrates MAPK-STAT3 signaling which in turn regulates the dynamic transition between 2 CAFs sub-populations, EMT-CAFs and proliferation-CAFs [48], rendering the IL6-TGFβ-EMT-CAFs cross-talk a valid therapeutic target. While targeting directly EMT via NOTCH or WNT has shown limited success in the clinic [49], microenvironment remodeling to reverse immunosuppression by inhibiting CXCL12 [50] or promoting T-cell infiltration [51] or function via engineered oncolytic adenovirus [52], has shown has shown promising results in reducing metastasis formation [53]. Additionally, we observed a positive correlation between gene expression of IL8, CXCL8, CXCR1 and CXCL10 and *Fn*/*Fusobacteriales* prevalence, corroborating findings from Casasanta assessing *Fn* in HCT116 CRC cells [54].

In conclusion, our analyses have identified a patient sub-population that has an unfavorable clinical outcome when their tumours exhibit mesenchymal traits and are highly positive with *Fn*/*Fusobacteriales* and pinpointed clinically-actionable host-specific vulnerabilities that suggest new treatments for these patients that extend beyond broad spectrum antibiotics.

## Materials and Methods

Detailed methods for the *in vitro* cell culture experiments and the study design, cohorts description and analysis steps are provided in the online supplementary materials and methods.

### Patient and public involvement statement

Patients or the public were not involved in the design, recruitment, conduct, reporting and dissemination of this research.

### Data availability

Processing and analysis code along with pathogen prevalence with corresponding clinical and molecular datasets for the Taxonomy and TCGA-COAD-READ cohorts included in this study will be made publicly available and archived upon publication at Zenodo (https://10.5281/zenodo.4019142). Pathogen prevalence will include *Fusobacterium nucleatum* load and *Fusobacteriales* relative abundance (along with higher resolution estimates at genus, family and species taxonomic rank) for the Taxonomy and TCGA-COAD-READ cohorts, respectively.

## Acknowledgments

We wish to thank the patients who kindly donated their samples and made this study possible. This study was supported by a grant from the Health Research Board and Science Foundation Ireland to JHMP (16/US/3301), a studentship to KS sponsored by The Northern Ireland Department for the Economy (NI DfE), and by funding from NI DfE (SFI-DEL 14/1A/2582; STL/5715/15). The results included here are in part based upon data generated by the TCGA Research Network: https://www.cancer.gov/tcga. We wish to acknowledge the Information Technology department at the Royal College of Surgeons in Ireland and the DJEI/DES/SFI/HEA Irish Centre for High-End Computing (ICHEC) for the provision of computational facilities and support.

