## Supplementary Materials and Methods for "Patients with mesenchymal tumours and high *Fusobacteriales* prevalence have worse prognosis in colorectal cancer (CRC)"

### In vitro experiments

#### Cell culture

HCT116 and HT29 cells were purchased as authenticated stocks from ATCC (Teddington, UK). HT29 cells were cultured in DMEM medium (ThermoFisher Scientific Inc.) supplemented with 10% fetal bovine serum (Invitrogen, Paisley, UK). HCT116 cells were cultured in McCoys 5A medium (ThermoFisher Scientific Inc.) supplemented with 10% fetal bovine serum (Invitrogen, Paisley, UK). Cell lines were screened for the presence of mycoplasma utilising MycoAlert Mycoplasma Detection Kit (Lonza) monthly and cultured for no more than 20 passages.

#### *Fn* culturing conditions

*Fusobacterium nucleatum* subsp. *nucleatum* strain 25586 was purchased from American Type Culture Collection (ATCC, Middlesex, UK). *Fn* was cultured at 37ºC under anaerobic conditions (DG250, Don Whitley Scientific, West Yorkshire, UK) in Fastidious Anaerobic Broth (Neogen, formerly Lab M, Scotland, UK).

#### Co-culture experiments

HT29 and HCT116 cells were co-cultured with *Fn* at a Multiplicity of Infection (MOI) of 10:1, 100:1 and 1000:1 under normal culturing conditions for the CRC cell lines.

#### Western Blotting

Western blotting analysis was carried out as previously described [1]. IκBα antibody (#9242) was supplied by Cell Signaling Technology (Danvers, MA) and β-actin (#A5316) was supplied by Sigma.

**NFκB activity assay**

Cells were co-transfected with NFκB luciferase reporter and Renilla constructs using X-tremeGENE HP (Promega, Madison, WI), as previously described [2]. Cells were lysed with Passive Lysis Buffer (Promega, Madison, WI) and Luciferase and Renilla activity assessed by luminescence using D-Luciferin and Colenterazine as substrates.

**Quantitative polymerase chain reaction (qPCR)**

RNA was extracted, according to manufacturer's instructions using the High Pure RNA Isolation kit (Roche, Burgess Hill, UK). The Transcriptor First Strand cDNA synthesis kit (Roche, Burgess Hill, UK) was utilized to synthesize cDNA, according to manufacturer's instructions. qPCR was performed on the LC480 light cycler, using Syber green, according to manufacturer's instructions. Primer sequences:

- **TNFα F**: CAG CCT CTT CTC CTT CCT GAT;
- **TNFα R**: GCC AGA GGG CTG ATT AGA GA;
- **β-tubulin F**: CGCAGAAGAGGAGGAGGATT;
- **β-tubulin R**: GAGGAAAGGGGCAGTTGAGT.

**Association between *Fusobacteriales* and *Fn* prevalence in tumour resections with host characteristics in CRC**

**Clinical cohorts**

In this study, we profiled *Fusobacteriales* and/or *Fn* in primary tumour tissue resections from n=645 CRC patients from an in-house (Taxonomy, [3]) and a public protected dataset (The

Cancer Genome Atlas, TCGA-COAD-READ). Demographic and clinical and pathological characteristics of the two cohorts are compared and contrasted in **Suppl. Table 1**, which was generated with the python package *TableOne* [5].

#### Taxonomy cohort

Stage II and III colorectal patients (n=156) from a multi-center study (St Vincent's Hospital, Dublin, IE; University Hospital Vall d'Hebron, Barcelona, ES; University of Aberdeen, UK; University of Florence, IT) were accrued, as previously described (Taxonomy cohort, [3]). The cohort collection was approved by the Medicine, Dentistry, and Biomedical Sciences School Ethics Committee (ref: 12/12v4), as previously described ([3]). In downstream analyses, we included patients with available gene expression profiling (Almac Xcel array, Almac Diagnostics, Craigavon, UK, GSE103479, [3]) and estimation of *Fn* load from resected tumour tissue (at least 50% tumour content) by qPCR (n=140). The primary outcome for the Taxonomy cohort was overall survival (OS), but disease-free survival (DFS) records were also available.

#### TCGA COAD-READ cohorts

Stage I to IV patients with cancer of the colon (COAD) or rectum (READ) accrued by The Cancer Genome Atlas (TCGA) network were considered for inclusion in the study (n=629). In downstream analyses, we included all patients (n=605) that i) where not listed as "Redacted" in the clinical metadata retrieved from Liu et al. [6]; and ii) had at least a high quality RNASeq experiment from primary tumour from which bacterial relative abundance could be estimated.

**Determination of *Fn* load and *Fusobacteriales* relative abundance in tumour resections of CRC patients**

**Taxonomy cohort**

*Fn* abundance was quantified through qPCR analysis from tumour DNA, performed on the Roche Light Cycler 480 Real Time PCR Instrument (Roche, Burgess Hill, UK), using Syber green, according to manufacturer's instructions. Each reaction contained 80 ng of genomic DNA which was assessed in duplicate, in 25 µl reactions. The abundance of *Fn* DNA in each tumour sample was normalised to the human reference gene Prostaglandin transporter (PGT) using the $2^{-\Delta Ct}$ method, where $\Delta Ct$ = Ct value for *Fn* – Ct value for PGT. Primer sequences:

- ***Fn* F**: CAACCATTACTTTAACTCTACCATGTTCA;
- ***Fn* R**: GTTGACTTTACAGAAGGAGATTATGTAAAAATC;
- **PGT F**: ATCCCCAAAGCACCTGGTTT;
- **PGT R**: AGAGGCCAAGATAGTCCTGGTAA.

**TCGA-COAD-READ cohort**

*Fusobacteriales* relative abundance in primary tumour specimens was estimated from RNASeq using a subtractive method implemented by the PathSeq pipeline (version 2, *PathSeqPipelineSpark* routine, [7]), powered by the Genome Analysis Toolkit engine (GATK, https://gatk.broadinstitute.org/, [9]) and the Apache Spark framework. Level 1 protected BAM sequencing files from RNASeq experiments for all TCGA-COAD-READ patients were accessed via the GDC Data Portal (https://portal.gdc.cancer.gov/) and served as input to the pipeline. Briefly, host reads (i.e. human) were filtered out and the remaining unmapped reads were aligned to microbial reads based on reference taxonomies for bacteria, fungi and viruses using a (default) min-clipped-read-length of 31. Host and microbe references files were retrieved from the GATK Resource Bundle (ftp://gsapubftp-anonymous@ftp.broadinstitute.org/bundle/pathseq/). We ran the PathSeq pipeline on n=698 patient samples of which n=644 were from tumour tissue. We restricted the analysis to samples which exceeded 10 million primary reads, resulting in n=630 high quality tumour samples for downstream analysis. Next, we collapsed microbial relative abundance from multiple samples and multiple tissue types (primary, recurrent and metastatic) of the same patient by mean. In downstream analyses, we included only patients with samples resected from primary tumours (n=605). We reported relative abundance for *Fusobacteriales* at the order, family, genus and species taxonomic rank as normalized score expressed as percentage of the total relative abundance of the bacterial kingdom. Some of the species, denoted by the suffix "_sp", such as *Fusobacterium_sp._CM1*, reported by PathSeq are actually sub-species/strains. This may lead to under-reporting the relative abundance of e.g. *Fusobacterium nucleatum* as it does not include the abundances of its sub-species/strains. To avoid this issue, we manually re-mapped sub-species/strains to their parent species by blasting their sequence in NCBI (https://www.ncbi.nlm.nih.gov/nuccore/). We performed the re-mapping only when the percentage of identity between the sub-species/strain and its parent species exceeded 97%, as indicated in the **Table** below. The majority of the sub-species/strains mapped to *Fn*.

**Table**. Sub-species/strain mapping to parent species.

| Sub-species/strain | Candidate parent species | Per. identity | Remapped parent species |
|---|---|---|---|
| Cetobacterium_sp._ZOR0034 | Cetobacterium_somerae | 100% | Cetobacterium_somerae |
| Cetobacterium_sp._ZWU0022 | Cetobacterium_somerae | 99.78% | Cetobacterium_somerae |
| Fusobacterium_sp._CM1 | Fusobacterium_nucleatum | 99.86% | Fusobacterium_nucleatum |
| Fusobacterium_sp._CM21 | Fusobacterium_nucleatum | 99.86% | Fusobacterium_nucleatum |
| Fusobacterium_sp._CM22 | Fusobacterium_nucleatum | 99.70% | Fusobacterium_nucleatum |
| Fusobacterium_sp._HMSC064B11 | Fusobacterium_nucleatum | 99.93% | Fusobacterium_nucleatum |
| Fusobacterium_sp._HMSC064B12 | Fusobacterium_nucleatum | 99.82% | Fusobacterium_nucleatum |
| Fusobacterium_sp._HMSC065F01 | Fusobacterium_nucleatum | 99.87% | Fusobacterium_nucleatum |
| Fusobacterium_sp._OBRC1 | Fusobacterium_nucleatum | 100% | Fusobacterium_nucleatum |
| Fusobacterium_sp._HMSC073F01 | Fusobacterium_varium | 100% | Fusobacterium_varium |
| Leptotrichia_sp._Marseille-P3007 | Leptotrichia_buccalis | 98.37% | Leptotrichia_buccalis |
| Leptotrichia_sp._oral_taxon_225 | Leptotrichia_trevisanii | 99.52% | Leptotrichia_trevisanii |
| Leptotrichia_sp._oral_taxon_879 | Leptotrichia_hongkongensis? | 96.86% | un-mapped |
| Leptotrichia_sp._oral_taxon_212 | Leptotrichia_hongkongensis? | 92.67% | un-mapped |
| Leptotrichia_sp._oral_taxon_847 | Leptotrichia_massiliensis? | 92.04% | un-mapped |
| Leptotrichia_sp._oral_taxon_215 | No candidate parent species found | un-mapped | |
| Fusobacterium_sp._oral_taxon_370 | Fusobacterium_nucleatum or Fusobacterium_periodonticum? | 95.33% for both | un-mapped |

### Gene expression analysis

For the Taxonomy cohort, transcriptomics data (Almac Xcel array, Almac Diagnostics, Craigavon, UK; GSE103479) were processed as previously described [3]. For the TCGA-COAD-READ cohort, level 4 batch-corrected and normalised gene expression profiles by RNASeq were retrieved from the TCGA PanCanAtlas data-freeze release (*EBPlusPlusAdjustPANCAN_IlluminaHiSeq_RNASeqV2.geneExp.tsv*) from https://gdc.cancer.gov/about-data/publications/pancanatlas).

### Transcriptomic-based signatures

We reviewed the literature and selected signatures encoding signaling pathways of interest including:

- **proliferation**: mean gene expression of BIRC5, CCNB1, CDC20, NUF2, CEP55, NDC80, MKI67, PTTG1, RRM2, TYMS, and UBE2C ([10]);
- **epithelial-to-mesenchymal transition (EMT)**: difference in gene expression of epythelial (CDH1, DSP, OCLN) and mesenchymal (VIM, CDH2, FOXC2, SNAI1, SNAI2, TWIST1, FN1, ITGB6, MMP2, MMP3, MMP9, SOX10, GCS) genes ([11]);
- **metastasis**: difference in gene expression of markers promoting (SNRPF, EIF4EL3, HNRPAB, DHPS, PTTG1, COL1A1, COL1A2, and LMNB1) and inhibiting (ACTG2, MYLK, MYH11, CNN1, HLA-DPB1, RUNX1, MT3, NR4A1, and RBM5) metastasis ([12]);
- **DNA damage**: mean gene expression of PRKDC, NEIL3, FANCD2, BRCA2, EXO1, XRCC2, RFC4, USP1, UBE2T, and FAAP24 ([13]);
- **WNT signaling**: mean gene expression of AC023512.1, APC, APC2, AXIN1, AXIN2, BTRC, CACYBP, CAMK2A, CAMK2B, CAMK2D, CAMK2G, CCND1, CCND2, CCND3, CER1, CHD8, CHP1, CHP2, CREBBP, CSNK1A1, CSNK1A1L, CSNK1E, CSNK2A1, CSNK2A2, CSNK2B, CTBP1, CTBP2, CTNNB1, CTNNBIP1, CUL1, CXXC4, DAAM1, DAAM2, DKK1, DKK2, DKK4, DVL1, DVL2, DVL3, EP300, FBXW11, FOSL1, FRAT1, FRAT2, FZD1, FZD10, FZD2, FZD3, FZD4, FZD5, FZD6, FZD7, FZD8, FZD9, GSK3B, JUN, LEF1, LRP5, LRP6, MAP3K7, MAPK10, MAPK8, MAPK9, MMP7, MYC, NFAT5, NFATC1, NFATC2, NFATC3, NFATC4, NKD1, NKD2, NLK, PLCB1, PLCB2, PLCB3, PLCB4, PORCN, PPARD, PPP2CA, PPP2CB, PPP2R1A, PPP2R1B, PPP2R5A, PPP2R5B, PPP2R5C, PPP2R5D, PPP2R5E, PPP3CA, PPP3CB, PPP3CC, PPP3R1, PPP3R2, PRICKLE1, PRICKLE2, PRKACA, PRKACB, PRKACG, PRKCA, PRKCB, PRKCG, PRKX, PSEN1, RAC1, RAC2, RAC3, RBX1, RHOA,

ROCK1, ROCK2, RUVBL1, SENP2, SFRP1, SFRP2, SFRP4, SFRP5, SIAH1, SKP1, SMAD2, SMAD3, SMAD4, SOX17, TBL1X, TBL1XR1, TBL1Y, TCF7, TCF7L1, TCF7L2, TP53, VANGL1, VANGL2, WIF1, WNT1, WNT10A, WNT10B, WNT11, WNT16, WNT2, WNT2B, WNT3, WNT3A, WNT4, WNT5A, WNT5B, WNT6, WNT7A, WNT7B, WNT8A (https://www.gsea-msigdb.org/gsea/msigdb/cards/KEGG_WNT_SIGNALING_PATHWAY);

- **Tumour Inflammation Signature (TIS)**: mean gene expression of CD276, HLA-DQA1, CD274, IDO1, HLA-DRB1, HLA-E, CMKLR1, PDCD1LG2, PSMB10, LAG3, CXCL9, STAT1, CD8A, CCL5, NKG7, TIGIT, CD27, and CXCR6 ([14]);
- **Cytolytic activity**: mean gene expression of GZMA, and PRF1 ([15]);
- **Interferon gamma (IFNγ)**: mean expression of IFNG, LAG3, CXCL9, and CD274 ([16]).

For both cohorts, we applied a robust scaling transformation (*sklearn.preprocessing.RobustScaler*) prior to computing the signatures. For the TCGA-COAD-READ cohort, gene expression profiles were quantile transformed (*output_distribution=normal*) prior to robust scaling.

### Characterization of the tumour microenvironment

Cell type composition was computationally deconvoluted from bulk tumour gene expression data using 2 methods: *Microenvironment Cell Populations-counter* (MCPcounter, [17]); and *quantification of the Tumor Immune contexture from human RNA-seq data* (quanTIseq, [18]). MCPcounter, implemented as R package, uses marker genes to estimate the abundance (in arbitrary units) of endothelial cells, fibroblasts and 8 immune cell types including T cells, CD8+ T cells, cytotoxic lymphocytes, B lineage, natural killer (NK) cells, monocytic lineage, myeloid dendritic cells and neutrophils. For the Taxonomy cohort, we computed MCPCounter estimates as previously reported [4] and we normalized the resulting scores using a robust scaler (*sklearn.preprocessing.RobustScaler*). For the TCGA-COAD-READ cohort, we applied a quantile-transform (*sklearn.preprocessing.QuantileTransformer* with optimal distribution set to normal) followed by robust scaling (*sklearn.preprocessing.RobustScaler*) prior to applying the MCPcounter algorithm. Cell type composition was further characterized by applying the quanTIseq pipeline (step 3 in *quanTIseq_pipeline.sh* from https://icbi.i-med.ac.at/software/quantiseq/doc/downloads/quanTIseq_pipeline.sh) to gene expression profiles of the Taxonomy ([4], flag set to account for the microarray nature of the data) or TCGA-COAD-READ cohort (*EBPlusPlusAdjustPANCAN_IlluminaHiSeq_RNASeqV2.geneExp.tsv*) without any additional pre-processing transformation. The quanTIseq algorithm uses a signature matrix to determine the fraction of tumour and stromal cells along with 10 immune cell types including non-regulatory CD4+ T cells, CD8+ T cells, regulatory T cells, dendritic cells, B cells, NK cells, neutrophils, monocytes, and classically- (M1) and alternatively- (M2) activated macrophages.

**Patients' classification into transcriptomic-based molecular subtypes**

Patients' tumour samples were classified according to the *Consensus Molecular Subtype* (CMS, [19]) and *Cancer Intrinsic Subtype* (CRIS, [20]).

Circa 20% of primary tumour samples cannot be classified as CMS1 to CMS4 and they are marked as "no label" (NOLBL, [19]). In order to maximize the number of patients with CMS assignments, patients were classified in CMS groups using the nearest prediction from the random forest (RF) classifier (R package *CMSclassifier*, https://github.com/Sage-

Bionetworks/CMSclassifier, [19]). For the Taxonomy cohort, we used the labels previously reported by McCorry et al. [4]. Similarly, for the TCGA-COAD-READ cohort, we retrieved the RF nearest prediction labels provided by Guinney et al. ([19], *cms_labels_public_all.txt* from synapse #: syn4978511). Additionally, we computed nearest prediction RF labels for the whole TCGA-COAD-READ cohort *de novo* to classify patients. We additionally included the CMS assignments for those patients that had not been subtyped as part of the Guinney et al. study. For both cohorts, subtype assignments mapping to multiple CMS classes were classified as indetermined and, thus, set to NOLBL.

Patients were subjected to CRIS subtyping and labelled as CRIS-A to CRIS-E or NOLBL (if Benjamini-Hochberg–corrected false discovery rate (BH.FDR) exceeded 0.2), as described in Isella et al. [20]. For the Taxonomy cohort, CRIS subtyping was performed using the nearest template prediction (NTP) classifier, available from GenePattern (https://genepattern.broadinstitute.org/gp/pages/login.jsf) as reported by McCorry et al. [4]. For the TCGA-COAD-READ cohort, we apply the CRIS subtyping to the whole TCGA-COAD-READ cohort. For the final CRIS assignments, we included either the labels provided from the Isella et al. publication [20] or the labels we computed *de novo* for patients that had not been subtyped as part of the original study.

### Unbiased and systematic analysis of human host associations with *Fusobacteriales* in the TCGA-COAD-READ cohort

#### Mutational status.

Genomic intra-tumour heterogeneity and mutational burden expressed as number of silent and non-silent mutations per Mb was retrieved from the supplementary materials of Thorsson et al. [21] and corresponding data-freeze (*mutation-load_updated.txt* from https://gdc.cancer.gov/about-data/publications/panimmune), respectively. Patients were classified as microsatellite stable (MSS) or unstable (MSI) using a cut-off of 0.4 applied to the MANTIS score retrieved from the supplementary materials of Bonneville et al. [22].

Somatic mutation data in Mutation Annotation Format (MAF, *mc3.v0.2.8.PUBLIC.maf.gz*) were retrieved from the TCGA PanCanAtlas data-freeze release (https://gdc.cancer.gov/about-data/publications/pancanatlas) and restricted to the subset of patients diagnosed with COAD-READ cancers. We used the *maftools* R package (version 2.2.10, [23]) to compute conversion changes (C>A, C>G, C>T, T>C, T>A, T>G) and the percentage of transitions (Ti) and transversions (Tv) from the MAF file.

For each patient and each gene, we extracted from the MAF file the number of detected mutational aberrations. As aberrations, we included frame shift deletions and insertions, in frame deletions and insertions, missense and nonsense mutations and splice sites and we excluded the following variants: 3' flank, 3' UTR, 5' flank, 5' UTR, Intron, RNA, silent and non-stop mutations.

Association between *Fusobacteriales* relative abundance (high vs. low using 75$^{th}$ percentile as cut-off) and mutational status (number of aberrations) was assessed with $\chi^2$ independence test. We restricted the analysis to genes with aberrations in at least 5% of patients (n=818 genes out of 21332, ~4%). We reported mod-log-likelihood P-values, adjusted for multiple comparisons with Benjamini-Hochberg FDR correction.

**Copy number alterations (CNAs)**

Copy number alterations (*broad.mit.edu_PANCAN_Genome_Wide_SNP_6_whitelisted.seg*) were retrieved from the TCGA PanCanAtlas data-freeze release (https://gdc.cancer.gov/about-data/publications/pancanatlas). Recurrent CNAs were identified in the TCGA PanCancer collection via The Genomic Identification of Significant Targets In Cancer (GISTIC, version 2, [24]) using a cut-off q-value of 0.25 and confidence threshold of 0.90 for peak boundaries (**Sup. Fig. 4**). A region was classified as amplification or deletion if the LogR was above or below the 0.1 threshold. Downstream analyses were restricted to patients from the TCGA-COAD-READ cohort with *Fusobacteriales* estimates (n=563). Copy number aberrations were visualised as a heatmap using the python package *cnvkit* (version 0.9.7, function *do_heatmap*), (**Fig. 3E**). Percentage of patients with aberrations at a given genomic position were visualised with the R package *copynumber* (version 1.26.0, function *plotFreq*, [25]), (**Sup. Fig. 5**). Differences in copy number aberrations at the cytoband level were computed by computing the difference in mean lesion frequency between patients with high vs. low *Fusobacteriales* relative abundance ($75^{th}$ percentile cut-off), (**Fig. 3F-G**).

**Aberrations in transcriptional and protein profiles**

A systematic screen was carried out to identify aberrations in transcriptional and protein profiles by *Fusobacteriales* relative abundance in patients of the TCGA-COAD-READ cohort. Association between *Fusobacteriales* relative abundance and either gene or protein expression was assessed by Spearman correlation (function *pairwise_corr* from the python package *pingouin*). P-values were adjusted for multiple comparisons for False Discovery Rate with Benjamini-Hochberg (function *pingouin.multicomp* from the python package *pingouin*). For transcriptional profiles, we restricted the analysis to the 5000 most variant genes. All available proteins were tested (n=189 proteins). Genes and proteins whose expression differed by *Fusobacteriales* relative abundance were put forward for pathway enrichment analyses carried out with the *gseapy* package (version 0.10.2) which provides a wrapper (function *gseapy.enrichr*) for *EnrichR* [26], (**Fig. 3 H-I, M-N** and **Sup. Fig. 6-7**).

**Exploration of putative mechanisms underlying differential impact of *Fn*/*Fusobacteriales* prevalence by tumour biology**

We fitted 2 logistic regression models to identify putative mechanisms underlying the differential impact of *Fn*/*Fusobacteriales* prevalence in mesenchymal vs. non-mesenchymal tumours. Specifically, we fitted:

- **model 1**: univariate logistic regression model (*Fusobacteriales ~ gene/signature*);
- **model 2**: logistic regression model with an interaction term for mesenchymal status (*Fusobacteriales ~ gene/signature * mesenchymal status*).

Patients were grouped into *Fusobacteriales*-low vs. high using the 75$^{th}$ percentile of *Fusobacteriales* relative abundance as cut-off. Selection of gene expression or signatures to include in model evaluation was hypothesis-driven. Tumour mesenchymal status was treated as binary (yes, no). Tumour were classed as mesenchymal if they were classified as CMS4 and/or CRIS-B based on transcriptomic assignments from the CMS [19] and/or CRIS [20] subtyping strategies. Logistic regression models were fitted using the function *statsmodels.formula.api.logit* from the python package (*statsmodels*).

### Statistical analysis.

#### Comparative analyses

For hypothesis-driven investigations, we visualized the association between either *Fn* or *Fusobacteriales* (order) relative abundance (high vs. low) with either split violin or mosaic plots drawn with the python packages *matplotlib* and *seaborn* for continuous and categorical clinical or molecular features, respectively. For hypothesis-driven analysis, we evaluated statistical significance by either non-parametric Kruskal-Wallis H-test for independent samples for continuous or categorical variables, respectively. Given the hypothesis-driven and exploratory nature of these analyses, the P-values were not adjusted for multiple comparisons. In contrast, in unbiased and systematic analyses (**Fig. 3**), P-values were adjusted for False Discovery Rate with Benjamini-Hochberg FDR correction (FDR-BH).

#### Outcome analysis.

As outcome endpoints, we evaluated both disease-free (DFS) and disease-specific (DSS) survival where we consider relapse or cancer-related death as event, respectively. For the Taxonomy cohort where the cause of death was not annotated, we assessed overall survival (OS, instead) and considered death by any cause as an event. We used Kaplan-Meier estimators and we fit univariate and interaction Cox proportional hazards regression models to evaluate survival by covariates. We assessed statistical significance with log-rank and likelihood ratio tests, respectively. Interaction Cox regression models were fitted to evaluate the cross-talk between pathogen prevalence (high vs. low using the 75$^{th}$ percentile as cut-off) and mesenchymal phenotypes (*mesenchymal*: either CMS4 and/or CRIS-B; vs. *non-mesenchymal*: neither CMS4 nor CRIS-B). For the Taxonomy cohort, we used *Fn* load as pathogen prevalence (**Fig. 5A-D** and

**Sup. Fig. 8**). For the TCGA-COAD-READ cohort, we used *Fusobacteriales* relative abundance as pathogen prevalence (**Fig. 5E-L** and **Sup. fig. 9**). In exploratory analysis, we additionally assessed the association between clinical outcome and pathogen relative abundance at higher taxonomic resolution (genus, family and species) for patients of the TCGA-COAD-READ cohort (**Fig. 5M** and **Sup. Fig. 10**).

Statistical analyses were performed in python (3.8.5), including the packages *tableone* (version 0.7.6, [5]), *statsmodels* (version 0.11.1), *pingouin* (version 0.3.8, [28]), *lifelines* (version 0.24.4, [29]). Statistical significance was set at $P<0.05$.
