## Supplementary Figures for "Patients with mesenchymal tumours and high *Fusobacteriales* prevalence have worse prognosis in colorectal cancer (CRC)"

**Supplementary Material**

**Supplementary Figures**

**Supplementary Figure 1.**

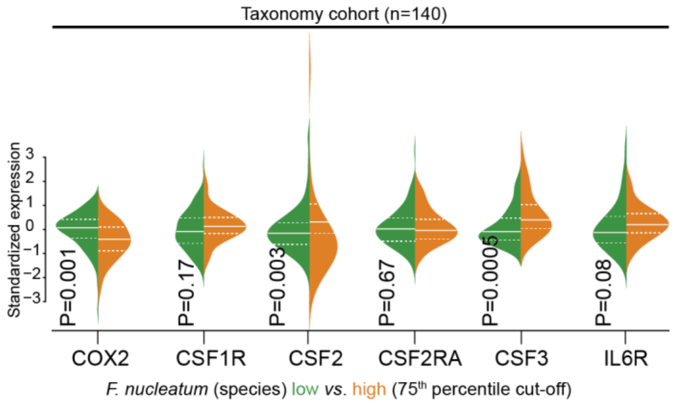


***Association between Fn load and inflammation signaling in the human host.***

Distribution of key player genes grouped by *Fn* (high vs. low, using the 75$^{th}$ percentile as cut-off) for patients of the Taxonomy cohort.

Median and lower (25$^{th}$) and upper (75$^{th}$) percentiles are indicated by white solid or dashed lines, respectively.

**Supplementary Figure 2.**

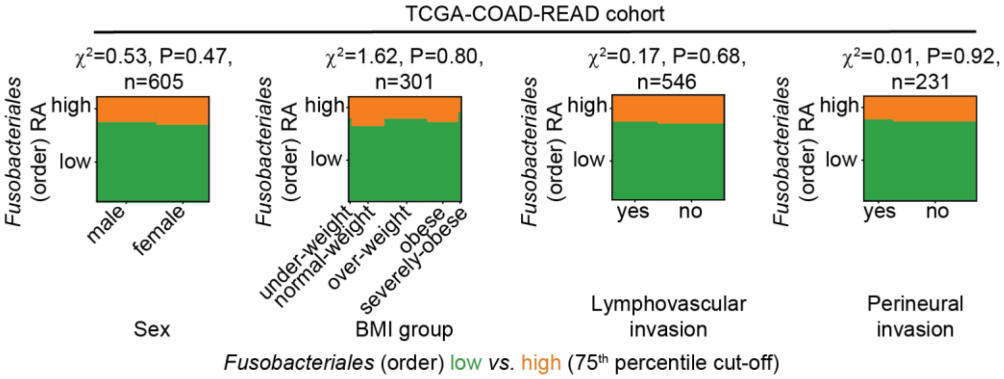


***Association between Fusobacteriales relative abundance and human host clinico-pathological features in the TCGA-COAD-READ cohort.***

Mosaic plots depicting the relationship between categorical clinico-pathological characteristics of the human host and *Fusobacteriales* RA. Patients were classified as *Fusobacteriales*-high or low using the 75$^{th}$ percentile as cut-off and indicated in orange and green, respectively. Statistical significance was evaluated with $\chi^2$ independence tests and the $\chi^2$ test statistic and the mod-log-likelihood P-values are reported.

***Abbreviations.*** BMI: body mass index.

**Supplementary Figure 3.**

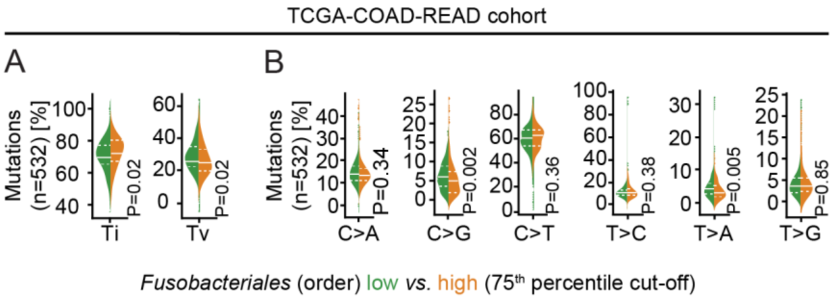


***Association between Fusobacteriales relative abundance and DNA substitution mutations in the patients of TCGA-COAD-READ cohort**.*

**A-B**. Distribution of transitions (Ti) and transversions (Tv) (**A)** and conversion changes (**B)** in patients of the TCGA-COAD-READ cohort classified as *Fusobacteriales*-high (in orange) or -low based on a 75$^{th}$ percentile cut-off. Median and lower (25$^{th}$) and upper (75$^{th}$) percentiles are indicated by white solid or dashed lines, respectively.

**Supplementary Figure 4.**

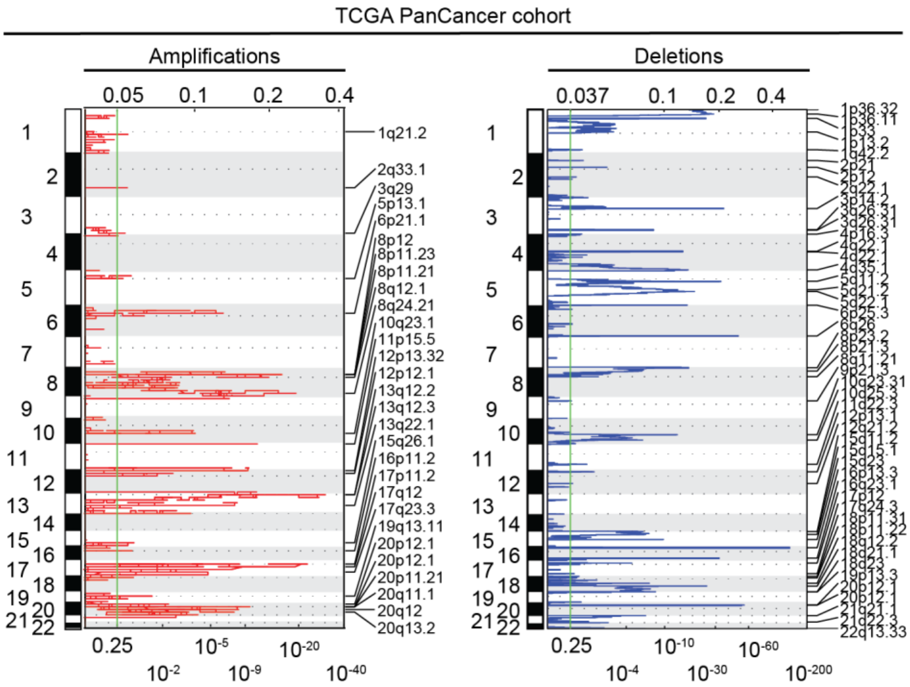


***Recurrent copy number alterations in patients of the TCGA PanCancer cohort.***

Amplifications (in red, left hand-side) and deletions (in blue, right hand-side) computed by GISTIC2 analysis to detect recurrent copy number alterations in the TCGA PanCancer cohort (n=9142).

Chromosome bands are indicated (y axis) and cytobands that reached statistical significance (as indicated by q-values) are shown.

**Supplementary Figure 5.**

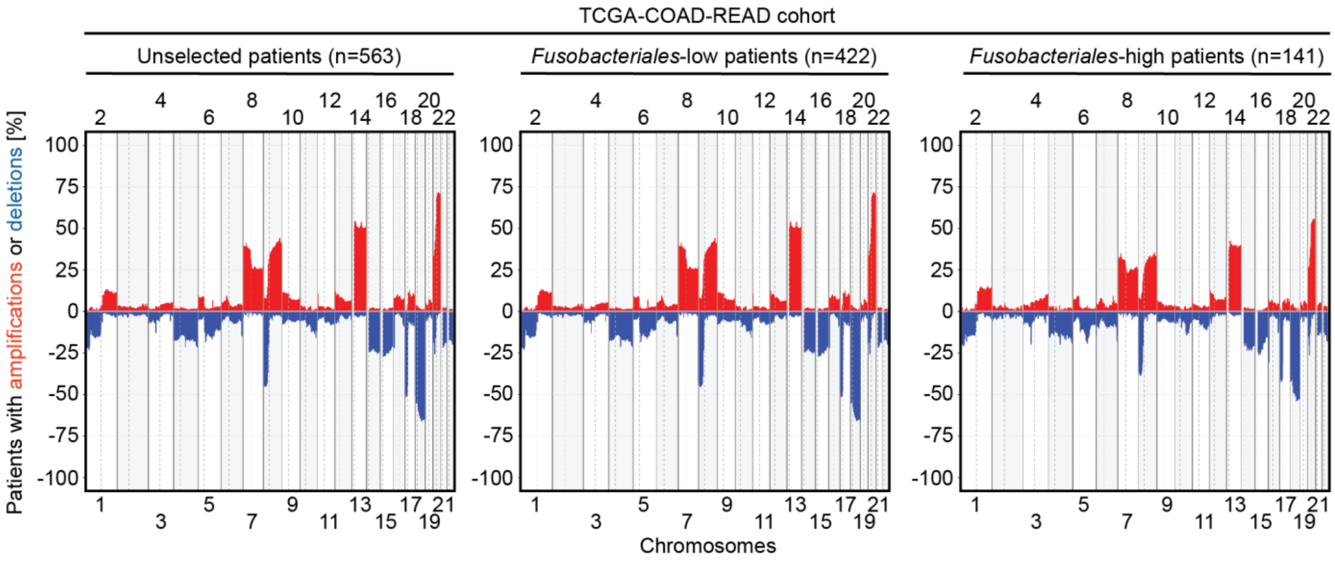


***Frequency of copy number alterations in patients of the TCGA-COAD-READ cohort.***

**A-C.** Frequency of copy number amplifications (in red) or deletions (in blue) by chromosome in patients of the TCGA-COAD-READ cohort. Figure's panels report frequency of occurrence in the whole unselected cohort (**A**) and in subgroups restricted to those patients with low- (**B**) or high- (**C**) *Fusobacteriales* relative abundance.

**Supplementary Figure 6.**

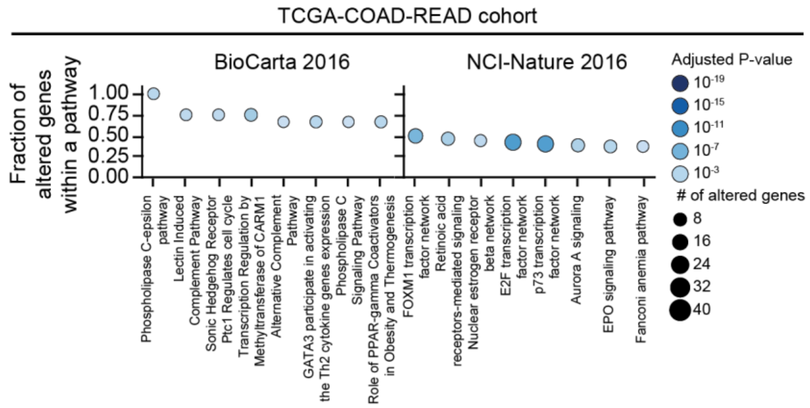


**Pathway enrichment analysis for genes differentially expressed by *Fusobacteriales* relative abundance in patients of the TCGA-COAD-READ cohort**.

Enrichment analysis on genes identified as differentially expressed by *Fusobacteriales* relative abundance in patients of the TCGA-COAD READ cohort. Analysis was performed with EnrichR querying the BioCarta (version 2016) and NCI-Nature (version 2016) pathway databases. The number of identified altered genes for each pathway is encoded by the marker size and the magnitude of the associated P-values is color-coded, as indicated in the legend.

**Supplementary Figure 7.**

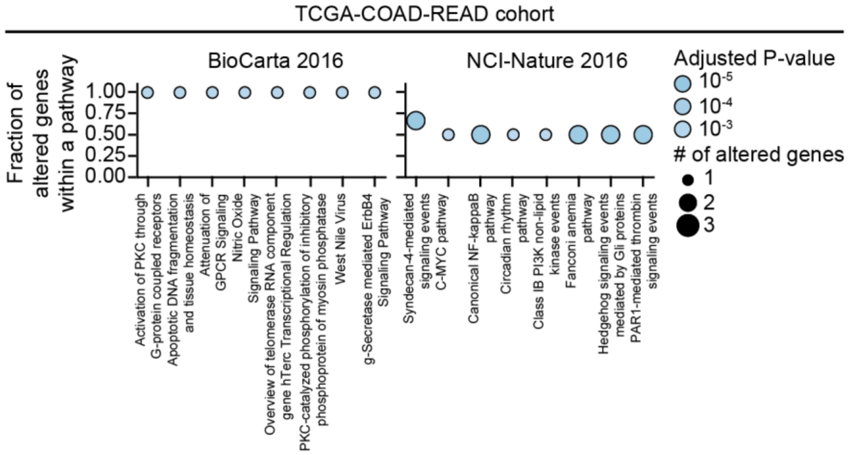


**Pathway enrichment analysis for proteins differentially expressed by *Fusobacteriales* relative abundance in patients of the TCGA-COAD-READ cohort**.

Enrichment analysis on proteins identified as differentially expressed by *Fusobacteriales* relative abundance in patients of the TCGA-COAD READ cohort. Analysis was performed with EnrichR querying the BioCarta (version 2016) and NCI-Nature (version 2016) pathway databases. The number of identified altered genes for each pathway is encoded by the marker size and the magnitude of the associated P-values is color-coded, as indicated in the legend.

**Supplementary Figure 8.**

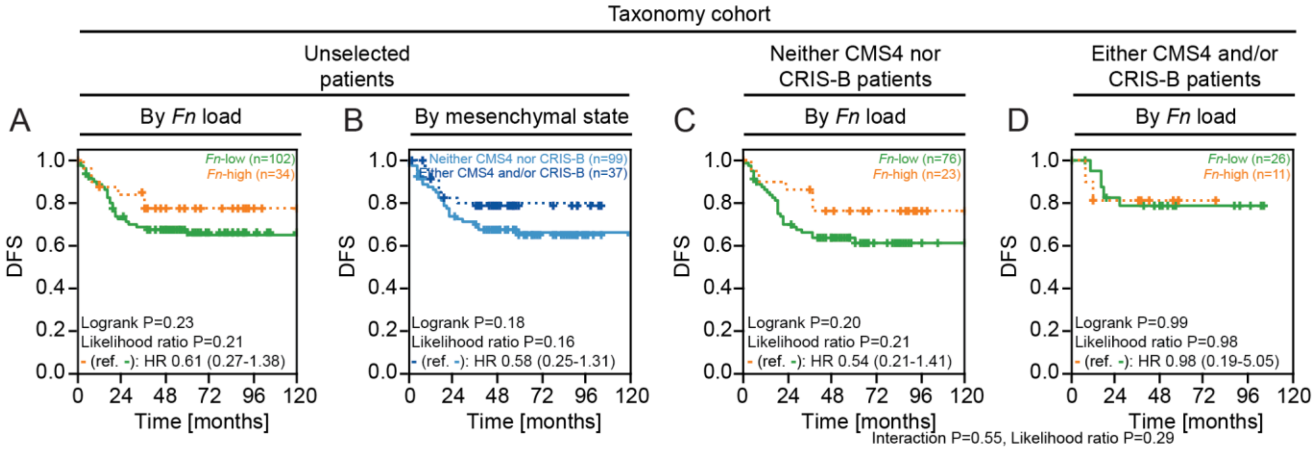


Kaplan-Meier plots comparing disease-free-survival (DFS) in patients of the Taxonomy cohort grouped by *Fn* load (**A**), mesenchymal status (**B**) and by *Fn* load within the non-mesenchymal and mesenchymal patients' subpopulations (**C-D**).

**Supplementary Figure 9.**

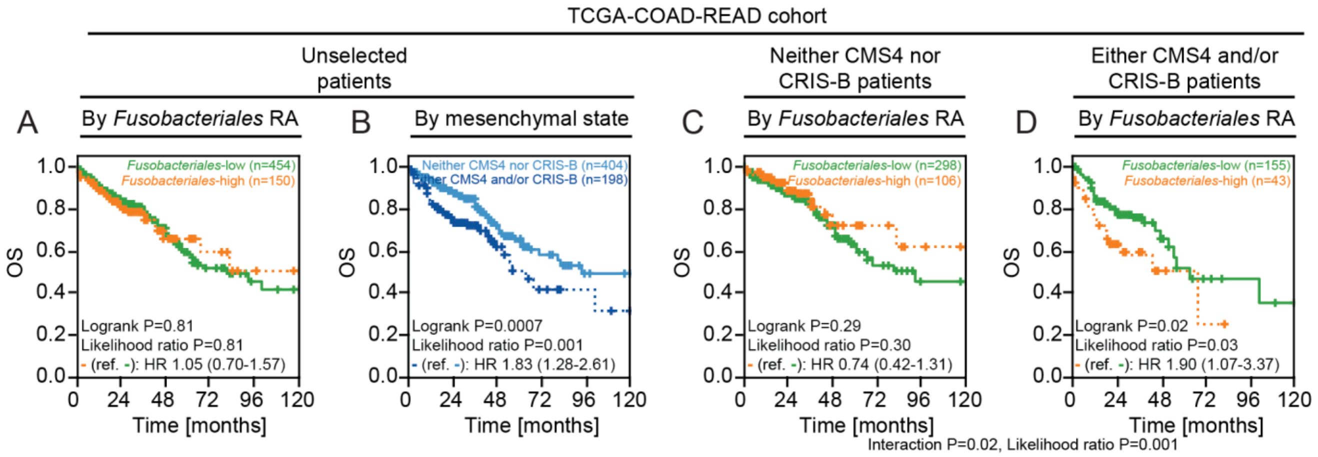


Kaplan-Meier plots comparing overall survival (OS) in patients of the TCGA-COAD-READ cohort grouped by **Fusobacteriales** RA (**A**), mesenchymal status (**B**) and by **Fusobacteriales** RA within the non-mesenchymal and mesenchymal patients' subpopulations (C-D).

**Supplementary Figure 10.**

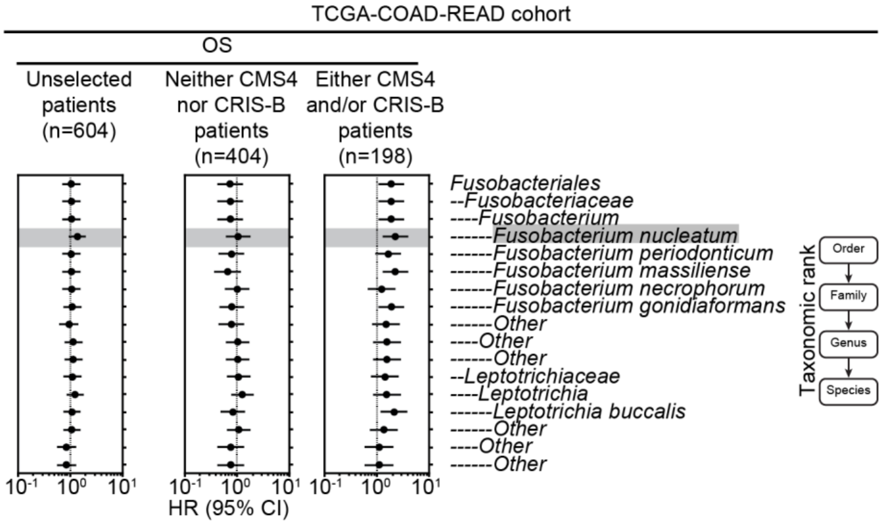


Cox regression models fitted on bacterium RA reported at the order, family, genus and species taxonomic ranks. For each taxonomic rank, patients were classified as low or high prevalence using the corresponding 75$^{th}$ percentile RA abundance as cut-off. Univariate Cox regression models were fitted when evaluating association between pathogen prevalence (high vs. low; reference low) at each taxonomic rank and OS in the whole unselected patient population (left panel). Cox regression models with an interaction term between pathogen prevalence (high vs. low; reference low) and mesenchymal status (mesenchymal, i.e. either CMS4 and/or CRIS-B, vs. non-mesenchymal, i. e. neither CMS4 nor CRIS-B) at each taxonomic rank and OS were fitted to evaluate differential impact of bacterium on clinical outcome by tumour biology (right panels).
