## Supplementary Tables Captions for "Patients with mesenchymal tumours and high *Fusobacteriales* prevalence have worse prognosis in colorectal cancer (CRC)"

### Captions for Supplementary Tables for

#### Supplementary Tables

##### Supplementary Table 1.

Clinico-pathological and demographic characteristics of the CRC patients of the Taxonomy and TCGA-COAD-READ cohorts.

##### Supplementary Table 2.

Association between mutational status and *Fusobacteriales* relative abundance in the TCGA-COAD-READ patients. Statistical significance was assessed by $\chi^2$ independence tests and $\chi^2$ statistics, unadjusted- and FDR-corrected mod-likelihood P-values are reported for each mutation that was either selected *a priori* or was found to be statistically significant altered when comparing *Fusobacteriales*-high vs. -low patients (75$^{th}$ percentile cut-off) of the TCGA-COAD-READ cohort.

##### Supplementary Table 3.

Association between recurrent copy number aberrations identified by GISTIC analysis when comparing *Fusobacteriales*-high vs. -low patients (75$^{th}$ percentile cut-off) of the TCGA-COAD-READ cohort.

##### Supplementary Table 4.

Association between gene expression profiles and *Fusobacteriales* relative abundance in the TCGA-COAD-READ patients. Statistical significance was assessed by Kruskal-Wallis H-tests and unadjusted- and FDR-corrected P-values are reported for each gene that was found to be statistically significant altered when comparing *Fusobacteriales*-high vs. -low patients ($75^{th}$ percentile cut-off) of the TCGA-COAD-READ cohort.

**Supplementary Table 5.**

Association between protein expression profiles and *Fusobacteriales* relative abundance in the TCGA-COAD-READ patients. Statistical significance was assessed by Kruskal-Wallis H-tests and unadjusted- and FDR-corrected P-values are reported for each protein that was found to be statistically significant altered when comparing *Fusobacteriales*-high vs. -low patients ($75^{th}$ percentile cut-off) of the TCGA-COAD-READ cohort.
