## Supplementary Table 1 for "Patients with mesenchymal tumours and high *Fusobacteriales* prevalence have worse prognosis in colorectal cancer (CRC)"

Clinico-pathological and demographic characteristics of the CRC patients of the Taxonomy and TCGA-COAD-READ cohorts.

| Variable | Level | Missing | Overall | In house Taxonomy | TCGA-COAD-READ | P-Value |
|---|---|---|---|---|---|---|
| n | | | 745 | 140 | 605 | |
| Stage, n (%) | I | 20 | 104 (14) | | 104 (18) | <0.001 |
| | II | | 294 (41) | 74 (53) | 220 (38) | |
| | III | | 241 (33) | 66 (47) | 175 (30) | |
| | IV | | 86 (12) | | 86 (15) | |
| T stage, n (%) | T1 | 1 | 22 (3) | 1 (1) | 21 (3) | <0.001 |
| | T2 | | 108 (15) | 6 (4) | 102 (17) | |
| | T3 | | 515 (69) | 101 (72) | 414 (69) | |
| | T4 | | 99 (13) | 32 (23) | 67 (11) | |
| N stage, n (%) | N0 | 3 | 417 (56) | 74 (53) | 343 (57) | 0.014 |
| | N1 | | 194 (26) | 49 (35) | 145 (24) | |
| | N2 | | 131 (18) | 17 (12) | 114 (19) | |
| M stage, n (%) | M0 | 72 | 588 (87) | 140 (100) | 448 (84) | <0.001 |
| | M1 | | 85 (13) | | 85 (16) | |
| Age, median [Q1,Q3] | | 1 | 68 [59,76] | 70 [62,78] | 68 [58,76] | 0.012 |
| Sex, n (%) | female | 0 | 341 (46) | 58 (41) | 283 (47) | 0.293 |
| | male | | 404 (54) | 82 (59) | 322 (53) | |
| Lymphovascular invasion, n (%) | no | 98 | 381 (59) | 57 (56) | 324 (59) | 0.664 |
| | yes | | 266 (41) | 44 (44) | 222 (41) | |
| Perineural invasion, n (%) | no | 413 | 262 (79) | 91 (90) | 171 (74) | 0.002 |
| | yes | | 70 (21) | 10 (10) | 60 (26) | |
| CMS subtyping, n (%) | CMS1 | 0 | 109 (15) | 28 (20) | 81 (13) | 0.002 |
| | CMS2 | | 300 (40) | 56 (40) | 244 (40) | |
| | CMS3 | | 105 (14) | 28 (20) | 77 (13) | |
| | CMS4 | | 170 (23) | 25 (18) | 145 (24) | |
| | NOLBL | | 61 (8) | 3 (2) | 58 (10) | |
| CRIS subtyping, n (%) | CRIS-A | 5 | 190 (26) | 36 (26) | 154 (26) | <0.001 |
| | CRIS-B | | 103 (14) | 20 (14) | 83 (14) | |
| | CRIS-C | | 187 (25) | 33 (24) | 154 (26) | |
| | CRIS-D | | 115 (16) | 17 (12) | 98 (16) | |
| | CRIS-E | | 132 (18) | 22 (16) | 110 (18) | |
| | NOLBL | | 13 (2) | 12 (9) | 1 (0) | |
